# Chronic Ethanol Exposure Produces Persistent Impairment in Cognitive Flexibility and Decision Signals in the Striatum

**DOI:** 10.1101/2024.03.10.584332

**Authors:** Yifeng Cheng, Robin Magnard, Angela J. Langdon, Daeyeol Lee, Patricia H. Janak

## Abstract

Lack of cognitive flexibility is a hallmark of substance use disorders and has been associated with drug-induced synaptic plasticity in the dorsomedial striatum (DMS). Yet the possible impact of altered plasticity on real-time striatal neural dynamics during decision-making is unclear. Here, we identified persistent impairments induced by chronic ethanol (EtOH) exposure on cognitive flexibility and striatal decision signals. After a substantial withdrawal period from prior EtOH vapor exposure, male, but not female, rats exhibited reduced adaptability and exploratory behavior during a dynamic decision-making task. Reinforcement learning models showed that prior EtOH exposure enhanced learning from rewards over omissions. Notably, neural signals in the DMS related to the decision outcome were enhanced, while those related to choice and choice-outcome conjunction were reduced, in EtOH-treated rats compared to the controls. These findings highlight the profound impact of chronic EtOH exposure on adaptive decision-making, pinpointing specific changes in striatal representations of actions and outcomes as underlying mechanisms for cognitive deficits.

## INTRODUCTION

Cognitive flexibility enhances survival by facilitating adaptation to changing environments thereby improving outcomes in an ever-changing world. The ability to make flexible decisions is substantially altered by chronic ethanol (EtOH) use and dependence across species, impairing goal-directed behavior while enhancing habitual or stimulus-oriented behavior^1–7^. Low cognitive flexibility may contribute to future escalation of substance use^8–10^, including EtOH intake in both rodents and primates^11–14^, and in heightening the risk of relapse to alcohol use in humans^15, 16^.

Cognitive flexibility, evaluated through reversal learning tasks^17, 18^, involves learning to make beneficial choices and adjusting those choices in the face of new action-outcome contingencies.

Although the cognitive and neural mechanisms underlying chronic EtOH-induced reversal deficits are not yet fully understood, studies in humans report persistent disruption of reversal learning during protracted withdrawal from EtOH drinking^19–22^. In rodents, EtOH exposure in adolescent subjects results in persistent cognitive deficits^23, 24^, but studies to date of chronic EtOH exposure and cognitive flexibility in adult rodents report relatively short-lasting (~< 2 weeks) deficits^25–28^, or no deficits^24, 29–31^. Thus, it remains unclear whether chronic EtOH exposure in adults causes a persistent deficit in flexible decision-making, despite evidence of such deficits in humans.

Corticostriatal circuitry is implicated in decision flexibility^32, 33^. In our previous studies, we found that chronic EtOH use induced maladaptive striatal synaptic plasticity^34–39^, particularly in the dorsomedial striatum (DMS), that persisted for several weeks^40^, in agreement with other recent work^3^. Thus, we hypothesize that repeated exposure to high concentrations of EtOH induces abnormal corticostriatal synaptic plasticity that disrupts striatal neural activity, leading to a persistent loss of cognitive flexibility.

In the current study, we confirmed that chronic EtOH-induced deficits after protracted withdrawal in adults are not apparent in standard two-bandit probabilistic reversal learning (PRL)^41, 42^. Thus, we devised a PRL task that requires rats to establish and update knowledge of reward probabilities under different levels of uncertainties. With this new task, we found EtOH-induced changes across multiple behavioral measures, including overall measures of the exploration-exploitation trade-off, as well as reversal deficits in highly uncertain environments. In addition, these deficits were found only in male, not female, rats. Using a reinforcement learning framework, we found that EtOH-exposed subjects showed specific alterations in differential value updating after receiving reward or no reward, providing a potential computational explanation for the observed deficits. Using *in vivo* electrophysiological recording within the DMS, we observed reduced encoding of choice signals alongside an enhanced encoding of outcome signals. Together, these findings reveal behavioral and neural mechanisms underlying the effects of chronic EtOH during protracted withdrawal and suggest that altered striatal processing of choice and outcome may contribute to EtOH-induced deficits in cognitive flexibility.

## RESULTS

### Protracted withdrawal from chronic EtOH exposure produces a distinct behavioral pattern in the dynamic PRL task

To investigate whether chronic EtOH exposure disrupts decision-making beyond the phase of acute withdrawal (i.e., >1 week), we used a well-validated chronic intermittent EtOH (CIE) vapor procedure to model EtOH dependence in rats^3, 43, 44^. Rats were trained on a standard probabilistic reversal learning (PRL) task and then underwent cycles of EtOH vapor or air exposure and withdrawal for four weeks (16 hrs/day, 5 days/wk), with group assignment balanced for PRL performance. Following 10-15 days of withdrawal, performance on the standard PRL task was reassessed (Figure S1A). This task required water-restricted rats to earn a drop (33 μl) of 10% sucrose water by choosing between left and right levers, which delivered a reward with either a 70% or a 10% probability, while the reward probabilities repeatedly switched within a session (Figure S1B). Previous studies have failed to find effects of the CIE procedure in adult rats on reversal performance^29, 30^. Indeed, during this standard PRL task, we did not observe any significant differences in reversal learning after block switches when comparing within-subject performance before and after the CIE exposure (Figures S1C-1G), nor between EtOH and air controls in either male or female rats (Figures S1C-1G). To further understand whether the CIE exposure could induce behavioral changes in this task, we employed a Support Vector Machine analysis (SVM)^45–47^ to decode group membership from a high-dimensional behavioral dataset consisting of 20 behavioral measures gathered during pre- and post-reversal phases in EtOH rats and their air controls (Figure S2A). We found that trained SVM models could not identify either group labels or sex labels in this PRL task above chance levels (Figure S2B). These findings indicate that rats did not express obvious EtOH-induced performance changes within the standard PRL task.

To further explore possible effects of chronic EtOH on cognitive flexibility, we next modified the task to present greater cognitive challenge^48, 49^. We introduced a three-block design with reward probabilities that varied in the contrast between left and right lever probabilities (80:10, 60:30, and 45:45; Figure 1C), converting the standard PRL task into a dynamic PRL task (dynaPRL). Consequently, when blocks transitioned from one to another, rats faced different amounts of uncertainty such that a larger change in the reward probabilities would be easier to detect and might facilitate adaptive changes in behavior (Figure 1C). Visual inspection of the trial-by-trial performance of individual rats in both air and EtOH groups revealed that subjects adjusted their choice behavior in response to block transitions within the dynaPRL task (Figure 1D). Applying the SVM model to the same behavioral features used for the analysis of behavioral data from the standard PRL task (Figure 1E), we found the model could decode group (air v.s. EtOH) and sex labels from the dynaPRL behavioral dataset with a prediction precision (63.29%) significantly higher than chance levels (25%; p <0.001). The trained SVM decoder almost never (<0.001% chance) misclassified male and female subjects (Figure 1G). More importantly, the decoding accuracy for EtOH treatment vs. air was significantly higher than chance for male but not female rats (Figure 1F). Using uniform Manifold Approximation Projection (UMAP), we found that the data were distinctly segregated for male EtOH and air rats in a 2D low-dimensional space with an accuracy (~70%), similar to the SVM decoding of group membership (Figure 1G).

**Figure 1.**
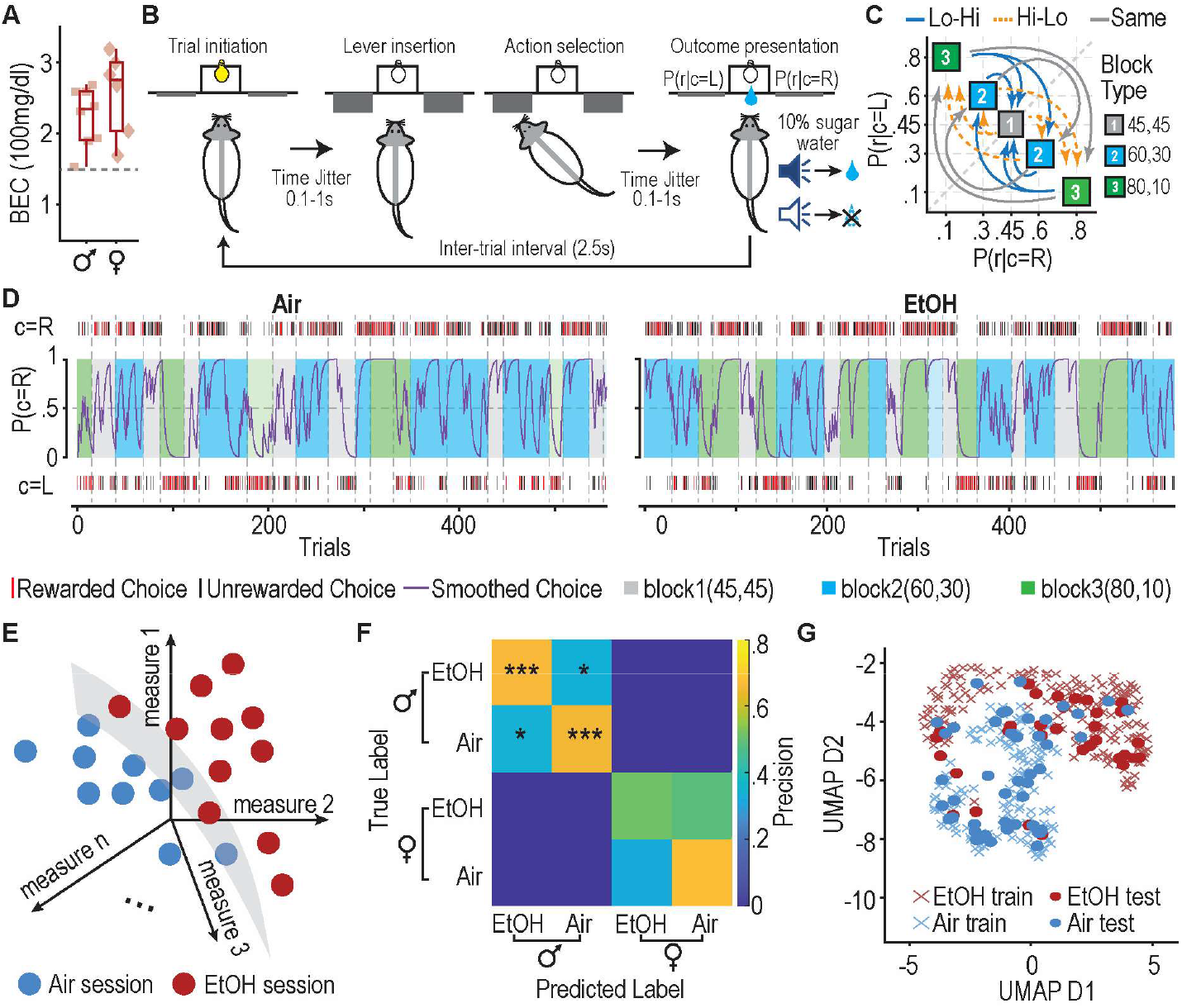
EtOH-exposed rats exhibit a distinct pattern of behavior in the dynamic probabilistic reversal learning task (dynaPRL) after prolonged withdrawal. (**A**) Blood ethanol (EtOH) concentrations (BEC) of male and female rats during EtOH vapor exposure. t(12) = −1.24, p = 0.239, t test. (**B**) Schematic of trial structure showing each behavioral stage in the dynaPRL task. Trials began with a magazine entry (trial initiation). After a random time interval between 100ms to 1s (time jitter), both levers were inserted. After the rat pressed one of the levers, the levers were withdrawn. Following another time jitter, an outcome was reported. A rewarded choice was indicated by a clicker sound, followed by sucrose, and an unrewarded choice was indicated by white noise. P(r|c=L) or p(r|c=R is the reward (r) probability given that the choice (c) is left (L) or right (R) side. (**C**) Choice reward probabilities for the dynaPRL task. The reward probabilities associated with each lever switched between 3 distinct blocks, drawing probabilities from three different sets, (0.8, 0.1), (0.6,0.3), and (0.45, 0.45). Based on the difference of reward probabilities between two levers before and after block transition, all block transitions can be categorized into three types: low to high (LH) challenge, high to low (HL) challenge, and same to same (no difference, ND), denoted as lines with distinct colors. (**D**) Two example rats from air (left) and EtOH (right) groups exhibit choice behavior that changes according to block switching. Choice raster, whether rewarded or not, are depicted with distinct colors. The letter ‘c’ denotes the side of choice, with ‘L’ for left and ‘R’ for right. The probability of choosing the right side, denoted as P(c=R), is calculated using a moving average across a window of 5 trials, thus providing a smoothed estimate of choice behavior. (**E**) This schematic illustrates how a hyperplane, depicted by the grey surface, differentiates between sessions from EtOH-exposed rats and control (air-exposed) rats, represented as dots, within the multidimensional space created by various behavioral measurements. This illustration represents our hypothesis that ethanol (EtOH)-exposed and air-exposed groups establish distinct behavioral patterns. (**F**) Confusion matrix for the SVM multiclass classifier aggregating from 1000 iterations in decoding both group and sex for n = 13 EtOH rats (8 male, 5 female) and 14 air controls (9 male, 5 female). Entries on the primary diagonal represent accurate predictions, and off-diagonal elements represent misclassifications. Classification precision was calculated based on the true positive rate/(true positive rate + false positive rate). *p < 0.05, ***p < 0.001, Monte Carlo significance tests. (**G**) A UMAP transformation of high dimensional data onto a 2D space.

Taken together, these results indicate that the impact of prior chronic EtOH exposure can be detected in male rats many weeks (>10 wks) after the last EtOH exposure through multivariate analysis of high dimensional behavioral measures during the more challenging dynaPRL decision-making task.

### Chronic EtOH exposure slows reversal learning and reduces exploration

The SVM analysis showed that behavior of male EtOH-exposed rats in the dynaPRL task differed from control rats. To understand the basis for this difference, we first examined how male EtOH and air rats adjust their choice behavior in response to changes in reward probabilities upon block switch. We analyzed the probability of selecting the action that was advantageous before the reversal. When a block switched from low uncertainty to high uncertainty (Lo-Hi) (Figure 2A), a two-way mixed-effect ANOVA revealed a significant interaction between group and trials (Figure 2B; p = 0.016). Bonferroni post-hoc tests showed that EtOH rats were significantly more likely to choose the previously preferred action in trials 2-5 after the block transition than rats in the air control group.

**Figure 2.**
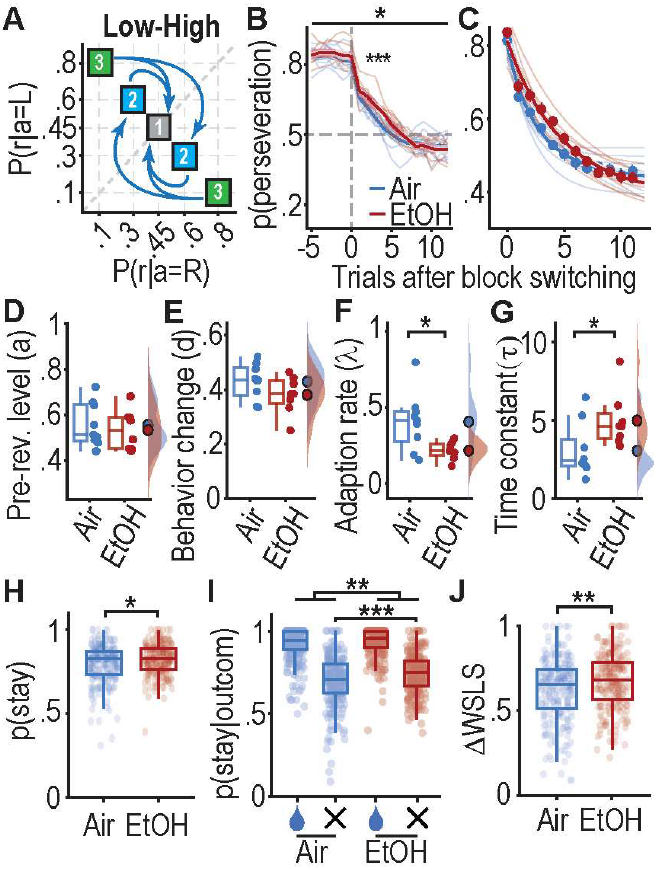
Protracted withdrawal from chronic EtOH exposure alters choice behaviors in highly ambiguous environments. (**A**) The diagram illustrates block switching in Low-to-High uncertainty transitions. **(B)** Probability of choosing the pre-switch preferred lever in Low-High type transitions. Bold lines represent the group average and faded lines represent individual rats. Vertical dashed line indicates the last trial before block switching (t=0). main effect of group, F(1, 16) = 1.708, p = 0.211; main effect of time, F(16,255) = 362.055, ***p = 1.2E-167; group x time interaction, F(16,255) = 1.925, *p = 0.016,, two-way mixed-effect ANOVA. *p < 0.05 for post hoc test with Bonferroni correction. (**C**) A single exponential decay model estimated the performance in learning new action-outcome contingency. Bold lines show the average model fit for probability of selecting an action based on the pre-switch contingencies. Dots depict the true mean data from B. Individual rat model estimates of choice probability are shown in faded lines. **(D-G)** The estimated parameters’ pre-reversal asymptotic value (pre-rev. level), a (D); magnitude of change in p(preservation) (behavior change), d (E); and adaptation rate, λ (F) for the single exponential decay model in C. The inverse of λ, τ, represents the time constant of rat’s learning in a new block (G). The box plot depicts the median, 25th, and 75th percentile of the parameter values for each group. Dots next to the box plot values from individual rats. The cloud plot on the right side of each panel depicts the data distribution. Dots on the cloud plot are the mean of the parameter values. t(15) = −0.498, p = 0.626 for D; t(15) = −1.522, p = 0.149 for E; t(15) = −2.704, *p = 0.016 for F; t(15) = 2.300, *p = 0.036 for G, two-sample independent t test with two-tails. n = 9 Air rats, 8 CIE rats for B-G. (**H**) Stay probability, the likelihood of repeating the same action on the current trial as the last trial, during ‘Lo-Hi’ transitions. z(464) = −2.358, *p = 0.018, Wilcoxon rank-sum test. (**I**) Stay probability given reward (water drop icon) or no reward (cross, X, symbol) on the last trial, during ‘Lo-Hi’ transitions. main effect of outcome, F(1, 463) = 1076.3, ***p = 7.3E-123; main effect of group, F(1, 463) = 10.625, **p = 0.001; outcome x group interaction, F(1, 463) = 4.5, *p = 0.035, two-way mixed-effect ANOVA. ***p < 0.001 post hoc test with Bonferroni correction. (**J**) Difference (Δ) between probability of stay given reward (win-stay, WS) and probability of shift given no reward (lose-shift, LS), ΔWSLS, during ‘Lo-Hi’ transitions. z(464) = −2.805, **p = 0.005, Wilcoxon rank-sum test. N = 241 sessions from 9 air rats, 229 sessions from 8 EtOH rats for B-J.

To quantify this further, we fit a single-exponential decay model to the probability of choosing the previously preferred action across the first 12 trials after the block transition (Figure 2C). While there was no difference in pre-reversal asymptotic value (*a)* or the magnitude of change in learning new action-outcome contingency after the reversal (*d*) (Figures 2D and 2E), we observed a lower adaptation rate (λ) in EtOH rats compared to the controls (Figure 2F; p = 0.016). Given that the inverse of λ (τ = 1/λ) corresponds to the time constant of the rat’s learning in a new block, this indicates that EtOH rats needed more experience to adapt to a new contingency in a highly ambiguous environment (Figure 2G; p = 0.036). Additional analyses applied to single sessions and individual rats revealed the same pattern of results (Figure S3A). By contrast, these effects were not observed during block transitions from high to low uncertainty (Hi-Lo), at the same uncertainty level before and after block transitions (same), or when all trial types were combined (Figures S3B-3D and S4A-S4F). Such findings suggest that the impact of EtOH on reversal learning dynamics is prominent only when facing a sudden transition to high uncertainty. To examine choice behavior later in a block, after rats are expected to have acquired the new contingencies, we focused on behavior >12 trials from block switch and assessed how much choice probability deviated from local reward rate yielded by the choice (‘matching behavior’)^50^. Male EtOH rats aligned their choices with local reward rates more tightly than same-sex control rats (Figure S5A–S5D).

To investigate strategies that might underlie the EtOH-induced choice behavior deficits, we analyzed behavioral metrics in two key phases, the early reversal learning phase, including trials 2 to 7 after transitioning to a new block, and the late phase of the current block, spanning the last 6 trials before block switch. We found that male EtOH rats were more likely to repeat the same choice in consecutive trials in the early phase following a switch, but not in the late phase after rats have adapted to the contingency change, as shown by measures of stay probabilities and the differences between win-stay and lose-shift probabilities (Figures 2H-2J and S6A-S6H). This behavior was sensitive to the receipt or omission of reward; EtOH rats exhibited higher stay probabilities after reward omission, but not reward receipt, during ‘Lo-Hi’ transitions (Figure 2I), resulting in a greater tendency for win-stay behavior and less tendency for lose-shift behavior (Figure 2J). We additionally found evidence that chronic EtOH exposure impacts deliberation processes, as revealed by alterations in latencies for trial initiation and choice during early reversal. Compared to air control subjects, EtOH rats were faster to initiate a new trial following a reward omission in the previous trial (Figures S7A–S7H) and had overall shorter latencies when making their choice, which was less modulated by the current action strategy (stay or switch) they held (Figures S8A-S8H). In contrast to males, female EtOH exposed rats did not show alterations in reversal behavior, matching behavior, win-stay behavior, or response latencies (Figures S4G-S4H, S5E-S5H, S6I-S6P, S7I-S7P, and S8I-S8P).

Collectively, these results demonstrate that chronic EtOH exposure significantly impacts outcome-driven action strategies, favoring a shift from switch to stay and enhancing matching behaviors in male rats, i.e., a tendency for greater exploitation and less exploration, resulting in a reduction in cognitive flexibility during a dynamic decision-making task.

### Chronic EtOH exposure alters value updating in reinforcement learning

To better understand the computational processes underlying the observed effects of EtOH exposure on decision-making, we applied several reinforcement learning (RL) models to the trial-by-trial choice and reward data. We found that a Q-learning model with differential learning rates (model 2, Q-DFLr) outperformed all other candidate models (Figure S9A). Q-DFLr has four distinct learning parameters controlling the updating of the value function: α^+^ and α^−^ for updating the value of chosen actions when the outcome is reward or no reward, respectively, and θ^+^ and θ^−^ for discounting the value of unchosen actions in an outcome-dependent fashion (Figure 3A). To assess the group-level differences between the EtOH group and the air control across RL parameters, we constructed a hierarchical model wherein the distributions of individual parameters for each rat are determined by the group-level hyperparameters. These hyperparameters encompass the means of the distributions for each parameter, the mean differences (δ) between air and EtOH groups for each parameter, and the variances of each parameter’s distribution. The posterior density of these hyperparameters was estimated through sampling using Hamiltonian Markov Chain Monte Carlo (HMC) methods. We found that distributions of group differences, δ, for α^+^ and θ^−^ substantially shifted towards the positive direction, indicating that the EtOH group has higher α^+^ and θ^−^ values than the air group (Figures 3B and 3C). We quantified the magnitude of evidence for group differences using directed Bayes Factor (dBF) and discovered that the likelihood of an increase in α^+^, the learning rate when rewarded, in the EtOH group was approximately 6-fold higher than a decrease (Figure 3B; dBF = 6.01). Similarly, for the EtOH group, the forgetting rate in the absence of reward, θ^−^, was roughly 8-fold more likely to increase than decrease (Figure 3C; dBF = 8.35). All other parameters showed small differences between the two groups (Figure S9B; all dBFs < 1).

**Figure 3.**
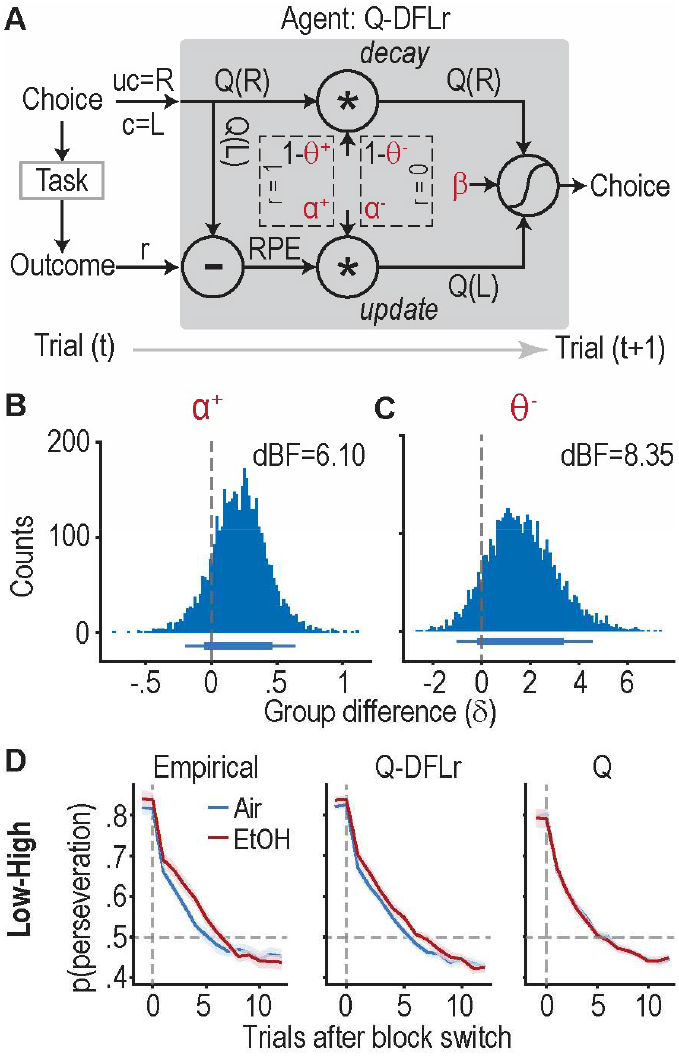
EtOH exposure enhances positive feedback-mediated learning rates and negative feedback-mediated forgetting rates in the reinforcement learning process. (**A**) Schematic representation of the value updating processes employed by the best-fitting reinforcement learning (RL) agent, Q-DFLr. In this example, an agent makes a left side choice at trial t, denoted as ‘c=L’, where ‘c’ stands for ‘choice’. ‘uc=R’ indicates the right lever was not selected, where ‘uc’ stands for ‘unchosen’. Upon making a left side choice, the task generates an outcome, ‘r’, where ‘1’ signifies reward, i.e., positive feedback, and ‘0’ signifies absence of reward, i.e., negative feedback. The left action value, Q(L)—or the chosen value—is updated based on the reward prediction error (RPE) using two distinct learning rates, α^+^ for reward (r=1) and α^−^ for non-reward (r=0) action outcomes. Conversely, the right action value, Q(R)—or the unchosen value—undergoes decay at two distinct forgetting rates, θ^+^ and θ^−^, depending on the action outcome, r. The updated action values are then adjusted by the inverse temperature parameter, β, to inform the agent’s next choice (t+1) via a softmax selection function. (**B**,**C**) HMC-sampled posterior densities of group difference, δ, between EtOH- and air-exposed rats in positive feedback-mediated learning rates (α^+^) in (B) and negative feedback-mediated forgetting rates (θ^−^) in (C), across group-level hyperparameters. Bottom horizontal lines represent the 80% and 95% highest density interval (HDI). dBF, directed Bayes factor. (**D**) Choice probability of the initially preferred lever in low-to-high transitions. Data on the left panel are empirical (reproduced from Figure 1G), while the right panel depicts simulated data using the best-fitting parameters.

We then used the best-fit parameters to simulate choice behavior within the original block structure used in the experiment and analyzed the probabilities of perseveration for each block transition type. We found that the simulated data partially recapitulated the EtOH-induced reversal deficits observed in the empirical data when transitioning from a low to a high uncertainty block, but not for other types of block transitions (Figures 3D and S9C). Collectively, these data suggest that chronic EtOH exposure selectively enhances both reward-based updates for selected actions and omission-based forgetting of unselected options, biasing action-selection towards previously reinforced choices. These results provide an algorithmic explanation for the changes in choice behavior we observed.

### Prior EtOH exposure alters the recruitment of DMS neurons representing choice and reward

Our behavioral analyses indicated that EtOH and air-control rats showed differences in action value updating upon outcome receipt, suggesting that EtOH might alter outcome processing. We therefore sought to examine the impact of EtOH on DMS neural dynamics during outcome presentation, defined as the 1000ms after the outcome (reward or reward omission) was revealed, in the dynaPRL task. We recorded a total of 524 and 326 single units in the DMS from EtOH and air groups, respectively (Figure S10). Using regression models, we first examined how neural activity in different time epochs throughout the trial was affected by the animal’s choice, the outcome of the choice, and their interaction, in the current trial (t) and previous trial (t-1). While choice-related signals were apparent in both groups throughout the trial (Figure S11A), a lower fraction of choice-modulated neurons was observed in EtOH rats than in control rats during the outcome presentation (Figures 4A, 4E, and S11A; p = 0.021 for 4E). Conversely, a larger portion of DMS neurons (~80%) was significantly modulated by the outcome in EtOH rats than in rats from air control group (Figures 4B, 4F, S11C; p = 0.018 for 4F). Further analysis into whether EtOH exposure affects the striatal representation of choice-outcome conjunction or the value of chosen actions during the outcome presentation phase revealed no significant group differences in these neural representations (Figures 4C, 4D, 4G, 4H, and S11E).

**Figure 4.**
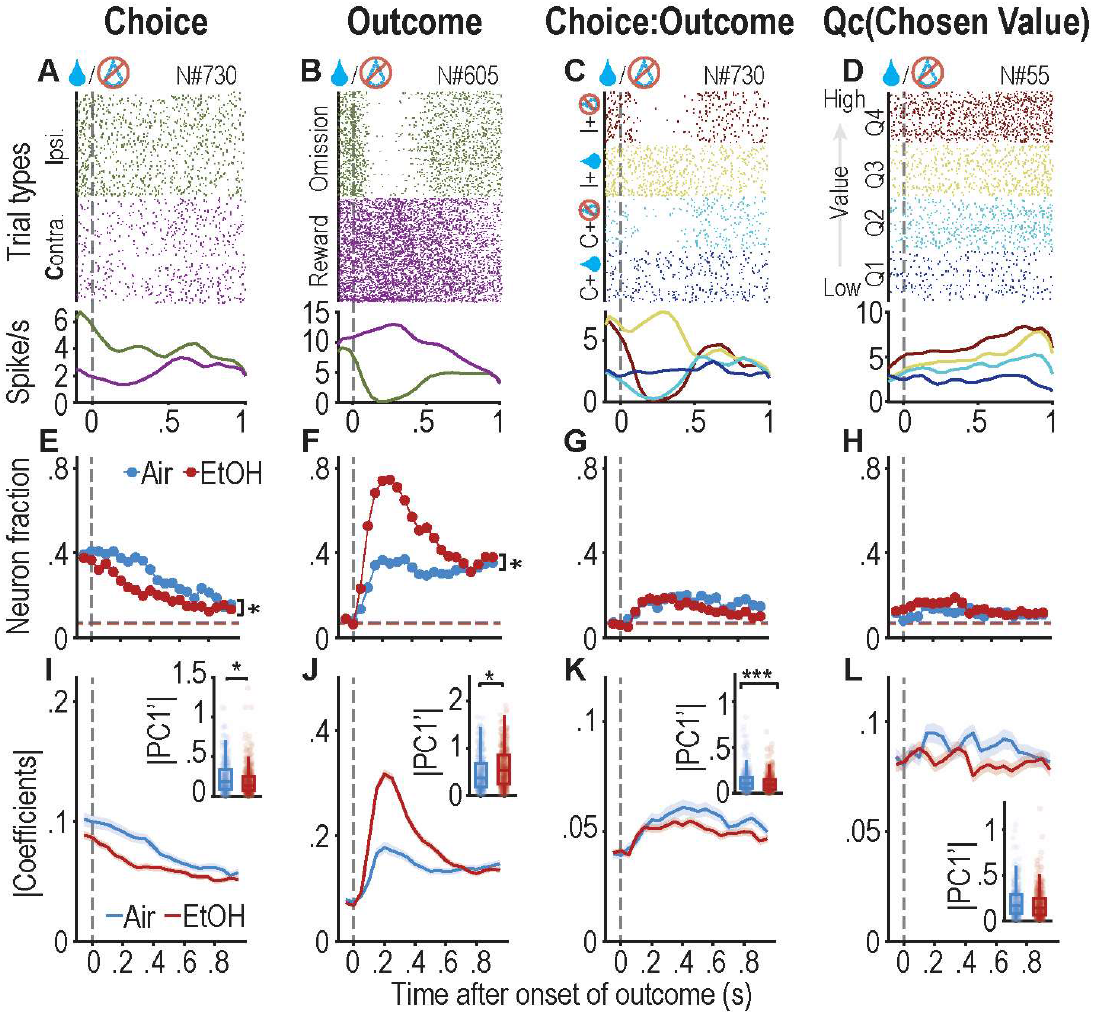
EtOH exposure alters encoding of decision variables in the outcome epoch. (**A**-**D**) Example DMS unit from the air group shows spike activity correlated with choice side (A), outcome (B), interaction of choice and outcome (labeled choice:outcome) (C), and the value of the chosen action, Qc (D). Trials are grouped by choice side as either ipsilateral (I) or contralateral (C) to the recording site in A; by outcome as rewarded (reward) or unrewarded in B; by contingency as ipsilateral choice without reward (‘I’ and no water drop icon), ipsilateral choice with reward (‘I’ and water drop icon), contralateral choice without reward (‘C’ and no water drop icon), and contralateral choice with reward (‘C’ and water drop icon) in C; and into quartiles of the chosen action value: Q1 (0-0.25), Q2 (0.25-0.5), Q3 (0.5-0.75), and Q4 (0.75-1) in D. At the top are spike raster plots. At the bottom are spike density functions generated by applying a Gaussian kernel (δ = 100 ms) to the corresponding spike trains from A-D. (**E**-**H**) Fraction of neurons with activity significantly modulated by the current trial’s choice (E), outcome (F), choice:outcome interaction (G), and Qc (H). *p = 0.021 main effect of group for E, logistic regression with Generalized Linear Mixed-Effect Model (GLMM); *p = 0.018 main effect of group for F, logistic regression GLMM; p = 0.47 main effect of group for H, logistic regression GLMM; p = 0.63 main effect of group for H, logistic regression GLMM. (**I**-**L**) The magnitude of the regression coefficient (|Coefficients|) for choice (I), outcome (J), choice:outcome interaction (K), and Qc (L) averaged across all neurons with the corresponding PC1 scores (|PC1’|) in the inset figures. ***p = 0.049 main effect of group for the inset figure in I, GLMM. *p = 0.041 main effect of group for the inset figure in J, GLMM. ***p = 0.0004 main effect of group for the inset figure in K, GLMM. p = 0.47 main effect of group for the inset figure in L, GLMM. n = 326 neurons from 9 air rats, 523 neurons from 8 EtOH rats.

The regression coefficients obtained from this analysis measure the impact of each decision variable on the neural firing rate during the period of outcome presentation. Given that the magnitude of these regression coefficients varied over time, Principal Component Analysis (PCA) was applied to the signed regression coefficients across time bins for each neuron. The magnitudes of the first principal component (PC1) scores were then utilized as an index of modulation strength over time. We found that choice and choice-outcome interaction signals have a significantly greater modulation strength on striatal firing activity from air rats compared to EtOH rats (Figures 4I and 4K; p = 0.049 and 0.0004, respectively), whereas outcome signals have a significantly greater modulatory effect on neural activity from EtOH rats compared to air controls (Figure 4J; p = 0.041). No difference was observed for the chosen value in modulating striatal firing (Figure 4L). While we found a significant impact of the previous trial’s outcome on neural activity within the trial, there was no evidence for group differences (Figures S11B, S11D, and S11F).

Collectively, these results demonstrate that a higher proportion of DMS neurons from EtOH rats encoded immediate outcome, whereas the signals related to current choice and action-outcome contingency were reduced relative to air controls.

### Action-selection value signals distinctly modulate DMS neuronal activity

Next, we evaluated whether EtOH impacted how value signals were encoded in the DMS during the action selection phase, defined as the time period from trial initiation until lever press. In particular, we sought to quantify the strength and time course of striatal activity related to two orthogonal combinations of the action values^52^ estimated by the best-fitting reinforcement learning model (model 2, Q-DFLr). The first variable corresponds to the difference between the two available action values, ΔQ, which is monotonically related to the probability of pressing the lever ipsilateral to the electrodes and is referred to as the policy. The second variable is the summation of the two available action values, ΣQ, and roughly corresponds to the so-called state value. We analyzed neural firing rates, aligned with either trial initiation, lever insertion, or lever press, as linear functions of ΔQ and ΣQ, with choice and chosen action value included as control variables in the regression analysis. We found that even several hundred milliseconds prior to trial initiation, a small (~15%) but significant proportion of striatal neurons from air control rats changed their spike rate in response to ΣQ (state value), and this state signal persisted until at least 200ms after trial initiation (Figures 5A and 5G). In contrast, the state value-sensitive neural population was much smaller in the EtOH rats (Figure 5G; p = 0.004). Thereafter, the proportion of neurons modulated by state value became similar in the two groups, decreasing close to chance levels during the lever insertion time epoch (Figures 5B and 5H) and increasing again after lever press (Figures 5C and 5I).

**Figure 5.**
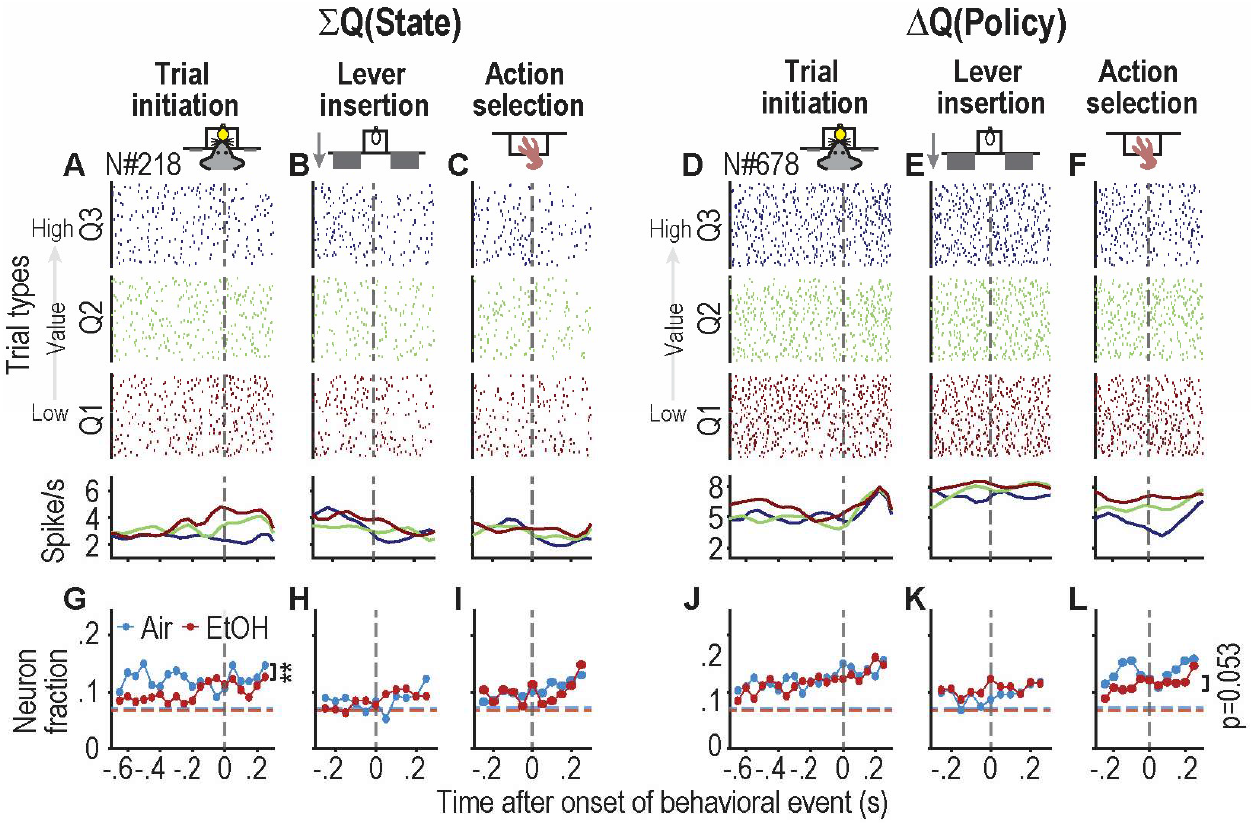
EtOH exposure alters encoding of decision variables in the action selection epoch. (**A**-**C**) Example DMS unit from the air group shows spike activity correlated with value of state, ΣQ, around trial initiation (A), lever insertion (B), and action selection (C). Trials are grouped by three quartiles of the state value: Q1 (0-0.33), Q3 (0.34-0.66), and Q3 (0.67-1) in A-C. At the top are spike raster plots. At the bottom are spike density functions generated by applying a Gaussian kernel (δ = 100 ms) to the corresponding spike trains from A-D. (**D**-**F**) Example DMS unit from the air group shows spike activity correlated with policy values, ΔQ, around trial initiation (D), lever insertion (E), and lever press (F). Trials are grouped by three quartiles of the policy value from low to high (bottom to top). Spike density functions at the bottom panels are estimated same as A-C. (**G**-**I**) Fraction of neurons with activity significantly modulated by state value, ΣQ, in the period of trial initiation (G), lever insertion (H), and lever press (I). **p = 0.004 main effect of group for G, logistic regression with Generalized Linear Mixed-Effect Model (GLMM). (**J**-**L**) Fraction of neurons with activity significantly modulated by state value, ΔQ, in the period of trial initiation (J), lever insertion (K), and lever press (L). p = 0.053 main effect of group for L, GLMM. figures. n = 326 neurons from 9 air rats, 523 neurons from 8 EtOH rats.

Additionally, a small proportion of neurons (~10%) encoded ΔQ (policy) starting before trial initiation; the fraction of these policy-coding neurons increased to about 15% until the completion of the lever press in air controls (Figures 5D, 5E, 5F, 5J, 5K, and 5L). There were no significant between-group differences in the proportion of neurons encoding policy during the trial epochs, although there was a tendency for a smaller proportion of policy-encoding striatal neurons in the EtOH rats than in the air controls when the spike rate was aligned to the lever press (Figure 5L; p = 0.053). These results suggest that EtOH differentially alters the neural representation of states selectively in the distal action selection phase (i.e., trial initiation) and possibly of action policies in the proximal action selection phase (i.e., lever press).

## DISCUSSION

In this study, we show that chronic EtOH vapor exposure during adulthood induces long-lasting (>10 weeks withdrawal) behavioral deficits in rats in a sex-specific manner. Behavioral data obtained during performance of a dynamic probabilistic reversal learning task (dynaPRL) could be used to predict group membership of EtOH-exposed vs air-exposed male subjects. Examination of the data revealed that cycles of high-dose EtOH exposure impaired later reversal learning under high uncertainty and led to an overall propensity to repeat a given choice, regardless of outcome, in male, but not female, rats. We employed a reinforcement learning model to elucidate underlying mechanisms for our observed behavioral deficits and found that the reward-mediated learning rate, which updates the value of chosen actions, and the reward omission-mediated forgetting rate that regulates the decay over time of the unchosen action value, were both increased in male rats exposed to chronic intermittent EtOH vapor. Lastly, EtOH exposure reconfigured firing patterns of dorsomedial striatum (DMS) neurons in male rats, augmenting neuronal sensitivity to the outcome while diminishing responsiveness to the choice, policy, and state signals. Our findings shed light on the cognitive and neural computational mechanisms underlying drug-induced persistent deficits in cognitive flexibility.

In humans, chronic EtOH abuse and withdrawal have been linked to marked neuropsychological impairments^22, 53–56^, with several studies highlighting cognitive deficits following moderate to extended periods of withdrawal^20, 21, 57–62^. Contrastingly, in rodents, while some long-lasting effects on goal-directed control^3, 63^ and working memory^64^ are noted, the evidence for impacts on reversal learning and attentional tasks has been inconsistent^24–31, 65^. This discrepancy between human and animal studies may stem from longer EtOH use or poly-substance use^66^ histories in humans and more complex cognitive demands of human tasks compared to rodent models. For example, the Wisconsin card sorting task involves complex rule shifts, unlike simpler rodent reversal learning tasks^33, 40–42, 67, 68^. Our findings from a standard probabilistic reversal learning (PRL) task showed no significant behavioral differences pre and post EtOH exposure, aligning with prior studies in both rodents^29, 30^ and humans^69^. However, by varying uncertainty in reward probabilities in an adjusted dynaPRL task^48, 49^ and hence enhancing its complexity and challenge, we uncovered more reliable EtOH-induced cognitive deficits.

When facing sudden cognitive challenge in the dynaPRL task, EtOH rats exhibited a robust deficit in reversal learning, i.e., a higher probability of repeating their choice, especially after reward omission in a high-uncertainty environment. Additionally, EtOH rats exhibited less deviation from matching their choices to the inferred reward schedule^51^. These findings suggest that EtOH rats might favor exploitation over exploration, in line with human studies^70^. Exploration-exploitation behaviors are modulated by novelty and uncertainty^71^. A plausible explanation for the reduction in exploration might be that EtOH pre-exposure diminishes outcome sensitivity. If true, a change in exploration across all other uncertainty conditions would be expected. However, this effect was observed only after transitioning from a low to high uncertainty environment. Therefore, another interpretation consistent with the current findings is that chronic EtOH exposure either decreases uncertainty-mediated exploration^72^ or promotes uncertainty avoidance following unrewarded choices, independent of learning^71^. Humans with EtOH use disorder display aberrant decision-making amidst high ambiguity in the Iowa Gambling Task^73, 74^ The intolerance of uncertainty has also been associated with risky EtOH use behavior^75^ and “hyperkatifeia”^76^. EtOH-exposed rats also exhibited shorter response times in the dynaPRL task across all block transitions (a change therefore independent of uncertainty) during protracted withdrawal. This observation aligns with increased impulsivity observed in humans with alcohol use disorder^77–79^. Thus, we identified multiple behavioral changes during protracted EtOH withdrawal in male rats.

We also found that male rats are more vulnerable to the negative impacts of chronic EtOH vapor exposure than female rats during the dynaPRL task. This is in line with previous literature that CIE vapor in adults rats impairs goal-directed behavior in males but not females^80^. Although Aguiree et al.^30^ showed that females are more vulnerable to chronic EtOH use in early reversal learning, this may be due to the fact that females consumed more EtOH than males in their study, whereas the vapor exposure method used in our study produced similar blood EtOH levels in both males and females, as in that of Barker and colleagues^80^. Estrogen potentially induces neuroprotective and anti-inflammatory effects^81, 82^ in chronic EtOH vapor-exposed female rats and facilitates recovery during the protracted withdrawal phase. However, this cannot fully explain the differences in behavior between males and females in the air control group. We speculate that the observed sex-related differences in response to uncertainty, particularly under stress, may stem from inherent variations in how males and females process uncertainty^83^. However, more studies are needed to understand how EtOH induces cognitive impairments in a sex-dependent manner.

Reinforcement learning theory provides a theoretical framework for describing the cognitive process of value-based decision-making^84^ and has been informative for the dissection of pathological decision processes in animal models of addiction^85^. Our observations in EtOH-dependent rats align well with findings from a human study utilizing a Q-learning model, which found that EtOH-dependent individuals have decreased sensitivity to reward omission and increased sensitivity to reward^86^. Similarly, in our study, a history of chronic EtOH exposure enhanced the updating of chosen action value upon receiving rewards, indicating heightened reward sensitivity. It also facilitated the decay of the unchosen action value when unrewarded, reflecting a diminished emphasis on negative feedback. As a result, EtOH exposure heightened the emphasis on selected actions regardless of their outcomes, leading to increased behavioral exploitation. This provides a computational mechanism that explains our observations of EtOH-induced behavioral perseveration.

Numerous studies support a critical role for the DMS in encoding decision-related variables^32, 52, 87–94^ and controlling action selection in value-based decision tasks^41, 95–100^. Expanding upon the findings from earlier research^91, 93^, we found that DMS neurons in rats from EtOH and air groups encode a preparatory choice signal at least few hundred milliseconds before trial initiation, peaking at the onset of the outcome report, and then diverging in EtOH and air treated rats, with reduced choice encoding in EtOH rats. During the same outcome period, encoding of the outcome is enhanced in EtOH rats. The observed aberrant attenuation of choice signals and augmentation of outcome signals during the outcome presentation period may disrupt contingency learning processes. Moreover, choice-outcome conjunction signals emerge post-reward onset and are impaired in rats with chronic EtOH pre-exposure, which may reflect an EtOH-induced deficit in representation of action-outcome contingency signals in the DMS. We also investigated the encoding of state value by DMS neurons. State values approximate how often an animal has been rewarded recently^52^, which is likely to be important environmental information for inferring the block type. Although each block has the same average reward probabilities, the local reward rate reflects the uncertainty level of the block (Figure S12). Our findings show suppressed state value representations at trial start in EtOH rats, which may disrupt the animal’s inference regarding the block transitions.

Our findings offer novel *in vivo* evidence that chronic EtOH exposure engenders enduring alterations in neurocomputational processes. It is possible the aberrant neural activity changes in the DMS might stem from EtOH-induced maladaptive plasticity in one or both of corticostriatal pathways and the nigrostriatal DA pathway. Prior studies found that chronic EtOH consumption and withdrawal induce aberrant orbito-striatal plasticity in the DMS^3, 37, 101, 102^. These orbitofrontal cortical inputs are proposed to encode various aspects of expected value^103–106^ and goal-directed choice and action-outcome associations^107–110^, respectively. Additionally, EtOH has been reported to disturb dopamine homeostasis in the striatum^111, 112^.

In summary, this study has unveiled specific and enduring behavioral and neural consequences of chronic EtOH exposure, providing new insights into mechanisms underlying cognitive impairments associated with EtOH addiction. Chronic EtOH consumption appears to distort neural computational processes, potentially interfering with the representation of decision-making signals, leading to cognitive inflexibility. These mechanisms may underlie some of the deficits observed in humans with alcohol use disorder. Importantly, behavioral and neural mechanisms for the changes we observed here after chronic EtOH might resemble those underlying loss of cognitive flexibility after chronic use of other drugs of abuse^113^ or other psychiatric disorders^114–116^.

## MATERIALS AND METHODS

### Animals

17 Male and 10 female Long-Evans (LE) rats (10 wks old upon arrival, Envigo) were singly housed. Rats were kept in a temperature- and humidity-controlled environment with a light:dark cycle of 12:12 h (lights on at 7:00 a.m.). Experiments were conducted during their light cycle. All animal care and experimental procedures were approved by the Johns Hopkins University Institutional Animal Care and Use Committee.

### Chronic intermittent EtOH vapor exposure (CIE)

Rats were exposed to four cycles of EtOH vapor (n = 8 male and 5 female) or air (n = 9 male and 5 female)^3, 44, 117, 118^. Each cycle consisted of 14 hrs of vapor exposure followed by a 10-hr withdrawal, repeated for 5 consecutive days. EtOH was volatilized by venting air through a heating flask containing 95% EtOH at a rate of 30-40 l/min. EtOH vapor was delivered to rats housed in Plexiglas chambers (La Jolla Alcohol Research, inc.). Blood was collected from the tip of tails in the middle and end of each round from all EtOH-exposed rats to assess blood EtOH concentrations (BECs) with an AM1 alcohol analyzer from Analox Inc. To control for the stress of the blood collection, we pinched the tip of tails for air control rats instead of cutting the tissue and collecting blood. Our procedure produced a mean BEC of 224.95 ± 14.51 (S.E.M) in male and 257.2 ± 23.79 (S.E.M) mg/dl in female rats without significant sex differences (Figure 1A, inset), which is in line with previous findings^3, 43, 44^.

### Probabilistic reversal learning (PRL) task

Rats undergoing behavioral testing were water-restricted to 90% of their *ad-libitum* weight. Behavioral experiments were performed in a soundproof customized modular operant chamber (Med Associates). The levers and a reward magazine were confined to the same wall of the chamber. Rats were initially trained to enter the magazine to collect a reward by delivering a 100 µl reward (10% w/v sucrose in tap water) at random times with an average interval of 60 s (RT60). Subsequently, rats were trained to initiate a trial by entering the magazine, which triggered the insertion of two levers into the chamber. Pressing a lever led to reward delivery (at this stage, both levers provided a reward with a 100% probability). Lastly, rats were trained on the reversal task, in which pressing one lever consistently delivered a reward (100% probability), and the other lever was not rewarded (0% probability). These action-outcome contingencies were reversed randomly 8 trials after the value of an exponential moving average over the past 8 choices (1 for correct and 0 for incorrect choice) crossed the 0.75 threshold. The block reversal probability was 10%. If the block was not switched within 20 trials of crossing the threshold, the computer forced the reversal. Thus, each rat had adequate trials to acquire current contingencies and could not predict the contingency reversal. One to three days after the deterministic reward schedule (100% and 0%) training, training in the final PRL task began. The reward probabilities for each lever were selected from 70% and 10%. Each trial in the final PRL task began with the illumination of the reward magazine. Upon a poke into the magazine (trial initiation), two levers flanking the left and right sides of the port were inserted after a variable delay (randomly selected between 100 ms to 1 s with 100 ms intervals). Animals were given 30-s to press a lever; failure to do so resulted in an invalid trial (<1% of trials for all rats) and was excluded from analysis. A lever press probabilistically resulted in either the CS− (0.5 s of white noise generated by MED Associated, ENV-225SM) with no reward delivery or the CS+ (two clicker sounds with a 0.1-s interval generated by MED Associated, ENV-135M) with ~33-μl reward delivery in the center magazine via a syringe pump (MED Associated, PHM-100). Each trial ended either at the end of the CS− presentation or after completing the reward collection (exit from the first magazine entry after reward delivery). Trials were separated by a constant 2.5-s intertrial interval. Each behavioral session lasted approximately two hours. All rats experienced 15 sessions of final PRL task prior to the vapor procedure and on average ~10.07 sessions of PRL task after the vapor procedure.

### Dynamic probabilistic reversal learning (dynaPRL) task

Water-restricted rats performed this task in the same operant chamber as in the original PRL task. The task structure was the same as PRL task except that reward probabilities assigned to each lever were drawn pseudorandomly from a set of blocks with paired probabilities [block1(neutral): (0.45, 0.45), block2(low contrast): (0.6, 0.3), block3(high contrast): (0.8, 0.1)] (Fig. 1F). Note that each block type had a distinct probability contrast (block1, 1:1; block2, 2:1; block3, 8:1), while maintaining the same averaged reward probability (0.45), and that the outcome uncertainty within a block varied with the degree of probability contrast across the two levers, i.e., high uncertainty in a low contrast block, etc. The lever outcome contingencies were randomly reversed to one of the three blocks every 15-30 trials, except when an animal made four or more consecutive incorrect choices within certain trial blocks (block2: 60,30 or block3: 80,10); such trials were excluded from trial progress. When switching blocks, the transition rules imposed two restrictions: 1) the position of the lever with the highest reward probability should not be the same in two consecutive blocks even if the reward probability contrast changes, and 2) Neutral blocks (block1) should not be repeated. Block transitions were categorized into three types based on the relative cognitive challenge level before and after each block transition. If the relative uncertainty was lower in the post-reversal block compared with the pre-reversal block, this was termed a ‘Hi-Lo’ transition (i.e., block2→block3, block1→block2, or block1→block3). If the relative uncertainty was higher in the post-reversal block compared with the pre-reversal block, this was termed a ‘Lo-Hi’ transition (i.e., block3→block2, or block2→block1, or block3→block1). If the relatively uncertainly was the same in the post-reversal block compared with the pre-reversal block while still shifting probabilities across the right and left levers, this was termed a ‘Same’ transition (i.e., block2→block2 or block3→block3) (Figure S2A). In addition, in this task, during non-neutral blocks (i.e., blocks 1 and 3) in 40% of the trials after the first post-reversal 12 trials, we probed the animal’s response to unexpected changes in reward size by doubling or halving the reward size (66 or 16.5 μl) compared to the remaining 60% of the trials. However, the probability of win-stay, latency of trial initiation, and action selection reaction time were not significantly modulated by reward size (Figure S13); thus, we will not focus on this feature in this manuscript. Rats were trained on the final dynaPRL task for at least 10 sessions prior to surgery.

### Electrode implantation

All rats were surgically implanted with custom-made drivable electrode arrays, with custom-designed 3D-printed pieces, sixteen insulated tungsten wires (50 um, A-M Systems), and two silver ground wires, soldered to two Plexon headstage connectors (Plexon Inc. TX) using standard stereotaxic procedures. Electrode arrays were placed unilaterally in the left or right DMS (AP: −0.1 mm; ML: 2.45 mm; DV: −4.7 mm from Bregma).

### Recording and Spike sorting

Following 1 week of recovery, rats were trained on the dynaPRL task again until they became accustomed to performing the task while their headstage was connected via a patch cable to the commutator located in the center of the chamber ceiling. Once their completed trials/session reached at least 80% of their pre-surgery average number of trials in a session (3-7 sessions), recording sessions began. Electrical signals and behavioral events were collected using Plexon multichannel neural recording systems (MAP system;) or Plexon OmniPlex systems, with a 40 kHz sampling rate. Neural recordings at the same depth in the DMS were obtained for multiple sessions; if multiple sessions from the exact recording location were included in the following analysis, the data from the same channel was included only once. After that, electrodes were lowered by approximately 160 μm, and recording resumed in the new location at least one day later.

Individual units were isolated offline using principal component analysis in Offline Sorter (Plexon) as described previously^92, 119–121^. Auto-correlograms, cross-correlograms, the distribution of inter-spike intervals, L-ratio, and isolation distance were used to ensure good isolation of single units. Spike and behavioral event timestamps were exported to MATLAB (MathWorks) using customized scripts for further analysis.

### Data Analysis and Statistics

#### Behavioral metrics

##### Reversal performance

Reversal performance was assessed by the probability of maintaining the same choice before the contingency change, denoted as *p(perseveration). p(perseveration)* was computed for each trial during the last 5 trials before and the first 12 trials after the contingency change. The *p(preservation)* was computed by the ratio of correct choices and the total number of blocks, except that if the pre-reversal block was a neutral block, in which neither side is correct, then a random side was selected as a pseudo-correct choice for the calculation. A single exponential decay function was fitted to the p(preservation) for the 1-12 trials after the contingency change,

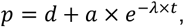

where *d, a*, and λ represent the pre-reversal asymptotic value, the magnitude of change in *p(perseveration)* after the reversal, and the learning speed, respectively. The single exponential decay function was defined by MATLAB function *fittype* and curve fitting employed MATLAB function *fit* with lower boundary (0, 0.5, 0) and upper boundary (3, 2, 1) for parameters d, a, and λ correspondingly. Averaged parameter values across 9 iterations were used for each subject and session.

##### Matching performance, stay probabilities, and latencies

To assess how animals allocate their responses (left and right lever presses) to the local (within a block) reward probabilities of each choice option after the learning period (first 12 trials), we calculated the *matching score*^50^ as a deviation of left response probability, *p(left)*, from the reward probability given select the left choice, *p(reward*|*choice=left)*. Deviation < 0 is undermatching, and > 0 is overmatching.

We measured the overall stay probability by calculating the likelihood of repeating the same choice as the last trial. Then, conditional stay probabilities were also calculated separately for the reward or unrewarded outcome in the last trial and denoted as *p(stay*|*win)* and *p(stay*|*lose)* correspondingly. The difference between win-stay and lose-shift was measured by ΔWSLS = *p(stay*|*win)* – *p(shift*|*lose)*.

The latencies (response times) measured the time interval between stimulus (center light illumination or lever insertion) and response or reaction (magazine entry or lever press) to that stimulus. As the distribution of latencies is highly skewed to the left (zero), a natural logarithm transformation was used to make the distribution of latencies more akin to normal distribution.

#### Classification analysis

##### Support vector machine (SVM)

SVM was employed to decode group membership (CIE vs. Air), sex label (male vs. female), and assessment phase (only for PRL task, pre-vapor vs. post-vapor) from the behavior dataset. SVM modeling was performed using the MATLAB function *fitclinear* for the classification of binary group membership and *fitecoc* for the classification of multi-subject labels. For both classification approaches, we used a total of 20 behavioral measures (columns) from each session consisting of reversal decay in post-reversal and matching score in pre-reversal, as well as 9 other behavioral measurements in pre- and post-block transitions, accordingly. The behavior dataset was randomly divided into two parts, a training set and a test set with a 90:10 ratio. Within the training set, a linear SVM model was optimized using 10-fold cross-validation. This approach involves dividing the training set into 10 equal parts, training the model on nine parts, and validating it on the tenth part. This process is repeated ten times, with each part used once as the validation set. The results from these iterations are used to fine-tune the model parameters. The optimized SVM model was then evaluated on a test set to assess its generalizability and predictive performance. This train-test process was repeated 1000 times. True positive (TP), true negative (TN), false positive (FP, type I error), and false negative (FN, type II error) were calculated based on actual predicted data labels. For the multilabel classification, precision [TP/(TP+FP)] was used to assess model performance in predicting each class. To determine whether the observed decoding accuracy for group membership and sex label was significantly higher than what would be expected by chance, we conducted Monte Carlo significance tests, and the p-value was computed as follows:

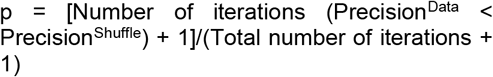

##### Uniform manifold approximation and projection (UMAP)

The manifold learning technique, UMAP, was employed to perform non-linear dimensional reduction of our behavior data matrix from above. We used the UMAP implementation provided in the MATLAB community *UMAP*^122^.

#### Reinforcement learning models

Reinforcement learning (RL) models assume that choices stem from the cumulative outcomes of actions accrued over numerous trials. We fitted the choice and outcome data to eight distinct RL models: 1, standard Q-learning model; 2, Q-learning model with differential learning rates (Q-DFLr); 3, differential forgetting RL model (DF-RL); 4, DF-RL model with inverse temperature β (DF-RLwβ); 5, forgetting RL model (F-RL); 6, F-RL model with inverse temperature β (F-RLwβ); 7, win-stay lose-shift model (WSLS); 8, biased random choice model (RD).

For models 1 through 4, choices are assumed to follow learned values for each option, Q, according to softmax function, in which the probability of choice k on trial t is:

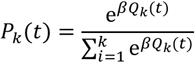

Here, β is the inverse temperature parameter that controls the level of stochasticity in the choice, with β=0 resulting in equal probability over all choice options. β was set to a constant 1 for models 4 and 6.

The value updating for each model is described below:

Model 1: Q-learning model,

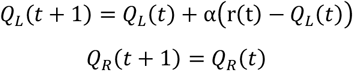

where *c*(*t*) is the choice side (left, L or right, R) in the current trial, *r*(*t*) is the outcome on that trial, and α, the learning rate, controls the update of the chosen action value according to the reward prediction error *r*(*t*) − *Q*_L_(*t*).

Model 2: Q-learning model with differential learning rates (Q-DFLr),

If *c*(t) = L and *r*(t) = 1,

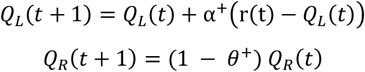

If *c*(t) = L and *r*(t) = 0,

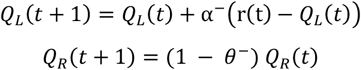

Here, value updates are governed by different learning rates for positive, α^+^, or omitted, α^−^, outcomes and the non-chosen option is decay by a forgetting rate, θ^+^ or θ^−^, for rewarded and unrewarded trials respectively.

Model 3 and 4: Differential forgetting RL (DF-RL) and DF-RL with (model 4) or without (model 3) free β,

If *c*(t) = L and *r*(t) = 1,

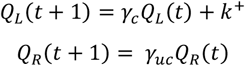

If *c*(t) = L and *r*(t) = 0,

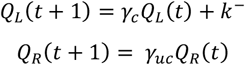

where κ^+^ and κ^−^ directly update the chosen action value in rewarded and unrewarded trials, respectively. γ_c_ and γ_uc_ are the forgetting factor in updating chosen and unchosen action values, respectively. Accordingly, forgetting rates θ_c_ and θ_uc_ in models 3 and 4 corresponds to 1-γ_c_ and 1-γ_uc_, respectively.

Model 5 and 6: Forgetting RL models (F-RL) and F-RL with (model 6) or without (model 5) free β,

If *c*(t) = L and *r*(t) = 1,

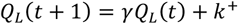

If *c*(t) = L and *r*(t) = 0,

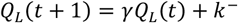

For the unchosen right side,

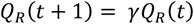

where κ^+^ and κ^−^ directly update the chosen action value in rewarded and unrewarded trials, respectively. γ is the forgetting factor in updating chosen and unchosen action values.

Models 7 and 8 directly describe the probability of each choice option on trial t, either subject to experienced outcomes (WSLS) or as a fixed proportion of choice.

Model 7: Win-stay lose-shift.

If *c*(t) = L and *r*(t) = 1,

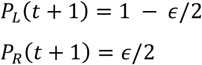

If *c*(t) = L and *r*(t) = 0,

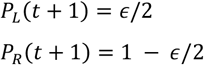

Where ϵ controls the randomness of the stay probability.

Model 8: Biased random choice.

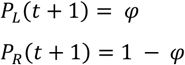

Where φ controls the bias of choice probability.

#### Hierarchical model fitting and model comparisons

We used the Matlab Stan interface (https://mc-stan.org/users/interfaces/matlab-stan) to construct and sample from hierarchical RL models, in which group-level hyperparameters govern individual rat parameters, to avoid overfitting to inter-rat noise ^123^. For a given model, we used Stan to draw consecutive samples from the posterior density of the model parameters using Hamiltonian Markov chain Monte Carlo (HMC), giving density estimates for the best fitting model parameters for the aggregate choice and outcome data across all sessions and rats. For each model, rat-level parameters were drawn from a group-level distribution with group mean, µ, group mean difference, δ, and variance σ. To quantify the evidence of group differences, the Directed Bayes Factor (dBF) was employed. This measure calculates the ratio of the proportion of the distribution of δ (group difference) that is above zero to the proportion that is below zero as in^115^. Given the difficulties in sampling from hierarchical RL models, parameters were transformed such that sample were drawn in an unconstrained space and later transformed into bounded values using the fast approximation of the cumulative normal distribution function, *phi_approx*^124^. We set normally distributed priors for all parameters at the rat level: N(0, 1) for all parameters, except β, which was set at N(0, 10), while group-level hyperparameters were given a prior normal prior distribution of N(0, 2) for µ and δ; and a gamma distribution of (2, 0.5) for σ. Hierarchical models for each of the candidate RL models were sampled for 2,000 iterations after 1,000 warm-up draws for four chains in parallel. All other settings adhered to the default configuration.

Predictive accuracy of each model was estimated using leave-one-out cross validation with Pareto-smoothed importance sampling (psis-loo) using the model log likelihood computed for each of the sampled parameters on the actual choice and outcome data^125^. For simulations of behavior, we took the mean of the posterior samples (i.e. over the 2000 post-warm up draws) for each rat-level model parameter as the best-fitting estimate for that individual subject. Using these best-fitting model parameters, we simulated choice behavior on the actual task structure. A reversal analysis, analogous to the one used on the empirical data, was applied to these simulated data to probe whether the model could recapitulate the key differences in choice behavior observed in the empirical data.

#### Multiple linear regression

To examine how neural spike activity was modulated by different behavioral and decision variables, we employed multiple linear regression models to fit spike rates of each unit as a linear function of various factors, such as rat’s choice, reward outcome, and estimated action values as in^32, 52^. All trials in each session other than those with choice omission were included in the analysis. To investigate the evolution of neural spike modulation within a temporal window surrounding key behavioral events, including lever insertion, action selection, and outcome presentation, we aligned the spikes of each neuron to the onset of these behavioral events. We then quantified the firing rates of each neuron using a sliding time bin approach. Specifically, we employed a 100-ms time bin with 50-ms steps or a 200-ms time bin with 100-ms steps. This analysis was conducted within predefined temporal windows relative to trial initiation, lever insertion, lever press, and outcome, presentation, which were set at −700 ms to 300 ms, −500 ms to 500 ms, −500 ms to 500 ms, and −100 ms to 1000 ms, respectively. To control for the temporal auto-correlation among spikes^126^, we added auto-regression (AR) terms as in Shin et al.^52^ for all of the following regression models. To investigate the influence of both current and preceding choices, outcomes, and their interaction on the spike activity of individual neurons, we employed a regression model to delineate the firing rate (*S*) as a function of choice direction (*C*; ipsi or contra-lateral to implant side), outcome (*O*; rewarded or unrewarded), and their interaction (*I*) in current trial *t* and last trial *t*-1, along with autoregressive components (*AR*), as the following

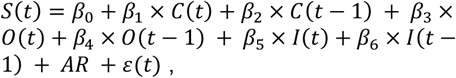

where β_0_ through β_6_ represent standardized regression coefficients, and the error term is ϵ(*t*). AR term was defined as follows:

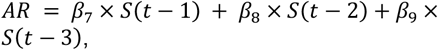

To examine how chosen value-related signals are represented by neurons at the time that choice and outcome are revealed, we used the following regression:

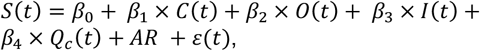

where Q_c_ is the chosen value estimated by the best fit RL model.

To investigate how different types of value signals are encoded in neuronal activity, we used a regression model as follows:

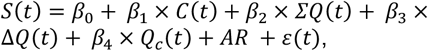

where *ΣQ*(*t*) = *Q*_ipsi_(*t*) + *Q*_contra_(*t*) and Δ*Q*(*t*) = *Q*_ipsi_(*t*) − *Q*_contra_(*t*), which can approximate state value and policy, respectively^32, 52^.

To capture the main variance of the coefficient across time bins, we ran a PCA on standardized regression coefficients. The magnitude of PC1 scores were used as a proxy of the overall modulation effect of decision factors on firing during the time window aligned to lever insertion, lever pressing, and outcome presentation.

#### Generalized linear mixed-effect models (GLMMs)

To understand the impact of EtOH treatment on the proportion of neural responses to specific decision variables described above, we employ a Generalized Linear Mixed-Effects Model (GLMM) to account for the random effects associated with individual animals and specific neurons within those animals. The model is described by the following Wikison notation:

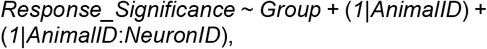

Where *Response_Significance* denotes the binary variable (e.g., significantly or not modulated) derived from the preceding multiple linear regression, and *Group* represents a fixed effect, such as an EtOH or Air group. The random effects are specified as *1*|*AnimalID* and *1*|*AnimalID*:*NeuronID*. The term *1*|*AnimalID* introduces a random intercept for each animal, allowing for the acknowledgment of variability in baseline response levels across animals. The term *1*|*AnimalID:NeuronID* adds another layer of random intercepts, this time to account for variability at the neuron level within each animal, acknowledging that responses may differ not just between animals but also among neurons within the same animal. To fit this model, we utilize the *fitglme* function in MATLAB with a ‘Binomial’ distribution and a ‘Logit’ link function.

To assess the influence of EtOH treatment on the overall modulational strength attributed to decision variables, we applied the following GLMM,

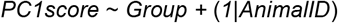

Here, *PC1score* represents the aggregate modulation effect derived from PCA on coefficients obtained from the aforementioned multiple linear regression analysis. This model was estimated using the *fitlme* function in MATLAB.

The p-value of the *Group* coefficient denotes the main effect of EtOH treatment.

### Other statistical analysis

Behavioral data were analyzed using paired *t*-tests, one-way ANOVA with repeated measures (one-way RM ANOVA), or mixed effect ANOVA (also known as two-way ANOVA with repeated measures, two-way RM ANOVA) followed by Bonferroni *post hoc* test, all conducted in MATLAB. A p-value < 0.05 was used as the criterion to determine statistical significance unless noted otherwise. All data are expressed as the mean ± standard error of the mean.

## Data availability

The datasets generated for this manuscript are available on request to. The code used in this manuscript is availble on github https://github.com/cyf203/CIE_flexibility_DMSrec ording upon publication.

## Ethics statement

All animal care and experimental procedures were approved by the Johns Hopkins University Institutional Animal Care and Use Committee and were conducted in accordance with the *National Research Council Guide for the Care and Use of Laboratory Animals*.

## Author contributions

Conceptualization: YC and PHJ. Reinforcement learning modeling: YC and AL. Analysis design for electrophysiology data: YC and DL. Other methodology, formal analysis and visualization: YC. Experiments and designs: YC and RM. Writing and discussion: YC, RM, AL, DL and PHJ. Supervision and funding: PHJ.

## FUNDING

This research was supported by R01AA026306 (PHJ) and R01DA035943 (PHJ) and in part by the Intramural Research Program of the NIMH under ZIAMH002983 (AJL).

---

**Figure S1.**
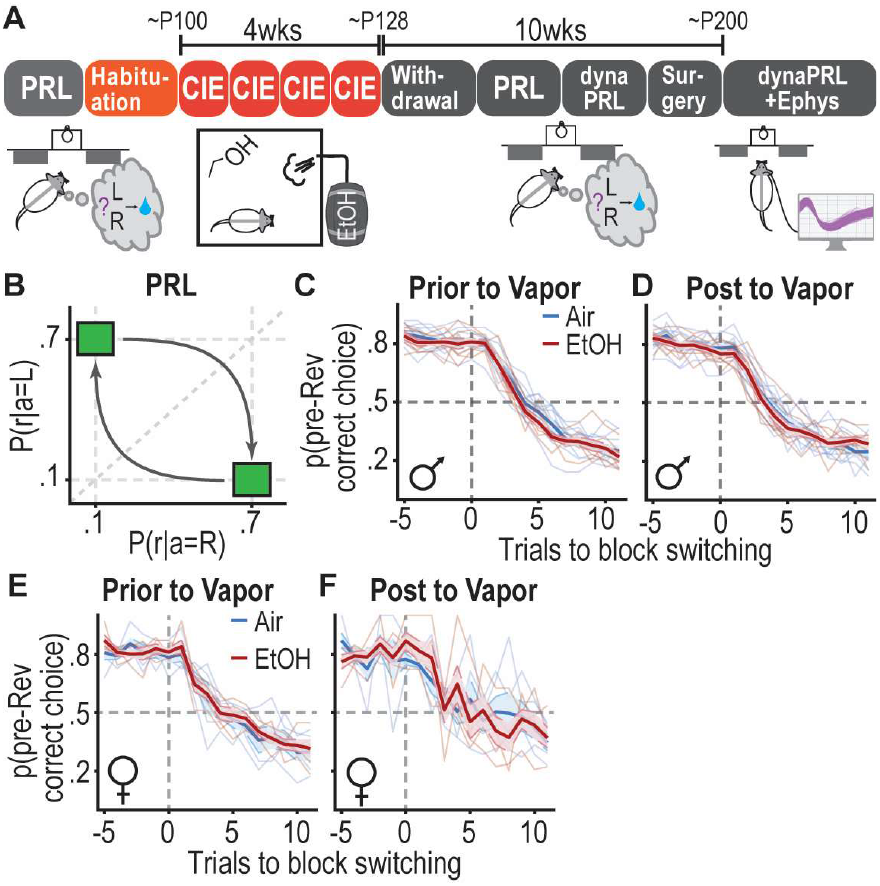
Protracted withdrawal from chronic ethanol exposure does not impair reversal learning in traditional probabilistic reversal learning task. (**A**) Experimental timeline: Rats underwent training in a simple probabilistic reversal learning (PRL) task to counterbalance their behaviors prior to 4-week chronic ethanol vapor exposure (CIE) procedure with a preceding habituation phase for acclimatization to ethanol vapor and experimental environment. Following 10-15 days of withdrawal from CIE, rats were retrained with the PRL task and then introduced to a new task, the dynamic probabilistic reversal learning (dynaPRL) task. 16-channel drivable electrodes were implanted in dorsomedial striatum of rats. After recovery and retraining, single-unit activity was recorded. (**B**) Choice reward probabilities for the PRL task. The reward probabilities associated with each lever switched drawing probability from a set (0.7, 0.1). (**C**,**D**) Probability of choosing pre-reversal preferred lever before (C) and after (D) CIE vapor in males. N = 9 male air rats, 8 male CIE rats. (**E**,**F**) Probability of choosing pre-reversal preferred lever before (E) and after (F) CIE vapor in females. N = 5 female Air rats, 5 female CIE rats. Bold lines represent the group average and faded lines represent individual rats. Vertical dashed line indicates the last trial before block switching (t=0) for D-G.

**Figure S2.**
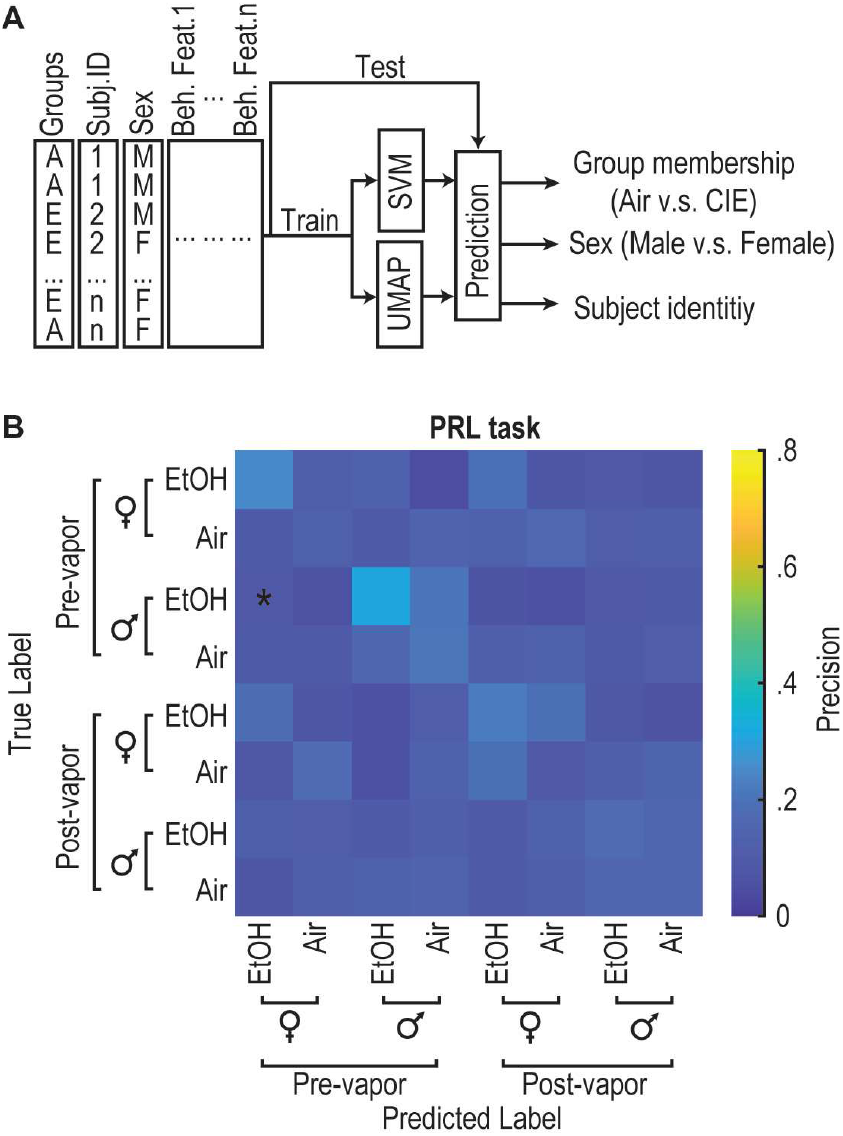
EtOH-exposed rats do not exhibit a distinct behavioral pattern in the standard PRL task. (**A**) The diagram illustrates the classification analysis process. (**B**) Confusion matrix for the SVM multiclass classifier aggregating from 1000 iterations in decoding group membership (EtOH vs. Air), sex label, and exposure conditions (pre- v.s. post-vapor) for N = 13 EtOH rats (8 male, 5 female) and 14 air controls (9 male, 5 female). Diagonal entries represent accurate predictions, while off-diagonal elements represent misclassifications. Classification precision is computed from true positive rate/(true positive rate + false positive rate). *p < 0.05 indicates statistical significance determined through Monte Carlo tests.

**Figure S3.**
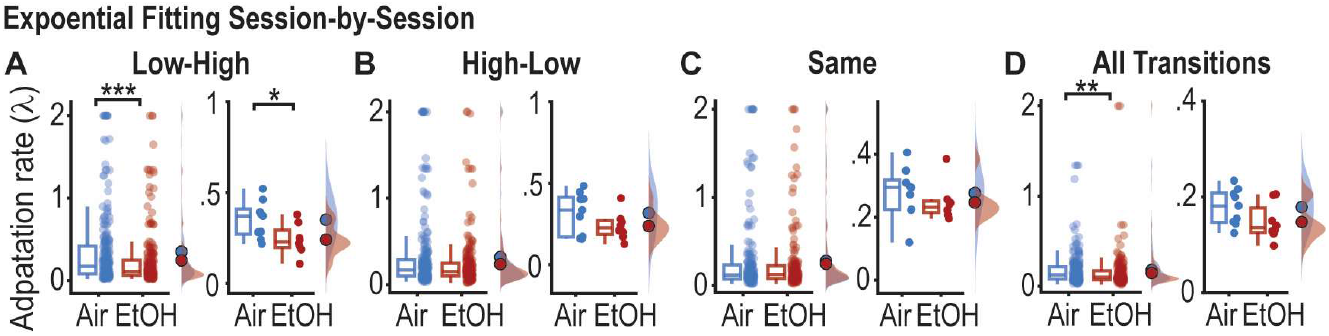
The adaptation rate (λ) is estimated across individual sessions in the dynaPRL task. (**A**-**D**) The adaptation rates (λ) in Low-High (A), High-Low (B), and Same (C) type transitions, as well as trials of all types of transitions (D) from individual sessions, are presented in the left panel in A-D. **p < 0.01, ***p < 0.001, Wilcoxon rank-sum test. The adaptation rates (λ) for each rat, averaged from individual sessions, are shown in the right panel of A-D. *p < 0.05, two-sample independent t test with two-tails. In the left panels, dots represent individual sessions. In the right panels, dots represent individual rats. The box plot depicts the median, 25th, and 75th percentile of the parameter values for each group. N = 241 sessions from 9 male air rats and 229 sessions from 8 male EtOH rats.

**Figure S4.**
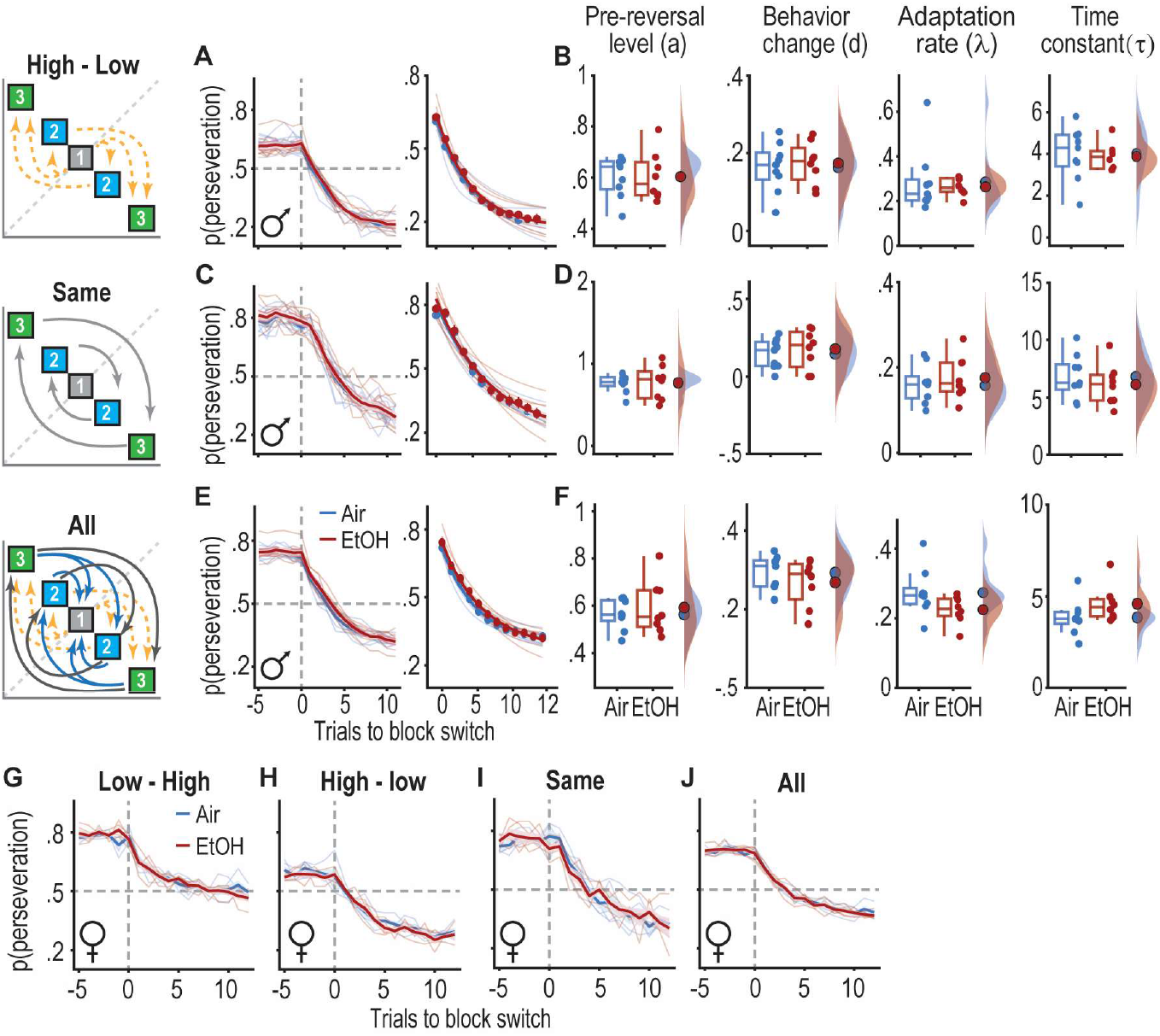
Prior EtOH exposure does not alter reversal learning under low uncertainty in the dynaPRL task. (**A-F**) The probability of selecting the preferred lever before the block switch (left panel) and the performance estimated by the single exponential decay model for learning new action-outcome contingency (right panel) during transitions with High-to-Low uncertainty (A), Same (C), or transitions aggregated across all types (E) in male rats. In the left panels, bold lines indicate the average model fit for the probability of selecting an action based on the pre-switch contingencies. In the right panels, dots represent the true mean data from A, C, and E, and faded lines illustrate model estimates of individual rat choice probability. The estimated parameters’ pre-reversal asymptotic value (pre-rev. level), a; magnitude of change in p(preservation) (behavior change), d, adaptation rate, λ, and time constant, τ, for the single exponential decay model during transitions with High-Low (B), Same (D), and transitions aggregated across all types (F) in males. Dots represent individual rats. The box plot depicts the median, 25th, and 75th percentile of the parameter values for each group. (**G**-**J**) Probability of selecting pre-switching preferred lever in transitions with Low-High (G), High-Low (H), and Same (I) type transitions as well as transitions aggregated across all types (J) in females. Bold lines indicate the average model fit for the probability of selecting an action based on the pre-switch contingencies. N = 13 EtOH rats (8 male, 5 female) and 14 air controls (9 male, 5 female).

**Figure S5.**
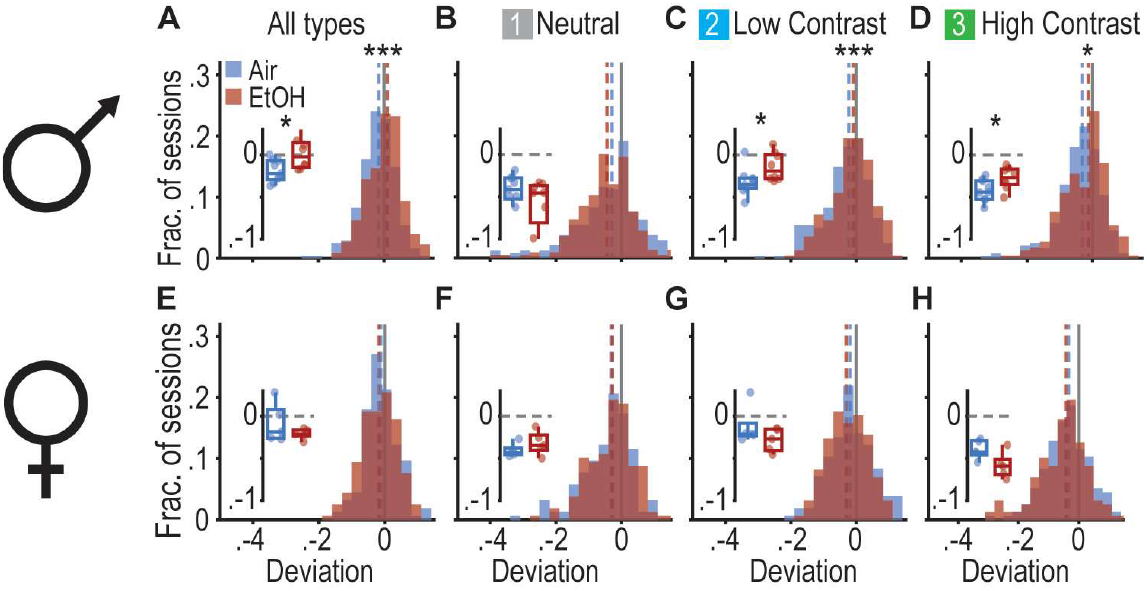
Prior EtOH exposure reduces deviation from matching choice behavior to local reward rate in males but not in females. (**A-D**) Histograms reveal higher matching scores (deviation of choice probability from the reward probability given the choice) in trials from blocks aggregating across all transition types (A), and low contrast (C; 60,30), as well as high contrast (D; 80,10) blocks, but not from neutral blocks (B; 45, 45), in EtOH rats compared to air controls. Vertical blue and red dashed lines represent the median deviation from matching, while the vertical grey solid lines represent perfect matching (0 deviation of choice probability to reward probability given the choice). The inset panels demonstrate the mean deviation score averaged for each EtOH rat and air control, with each dot representing a rat. The horizontal dashed line marks the perfect matching. (**E**-**H**) Similar to A-D, histograms show no difference in matching scores in all (E), neutral (F), low contrast (G), and high contrast (H) type blocks. N = 13 EtOH rats (8 male, 5 female) and 14 air controls (9 male, 5 female).

**Figure S6.**
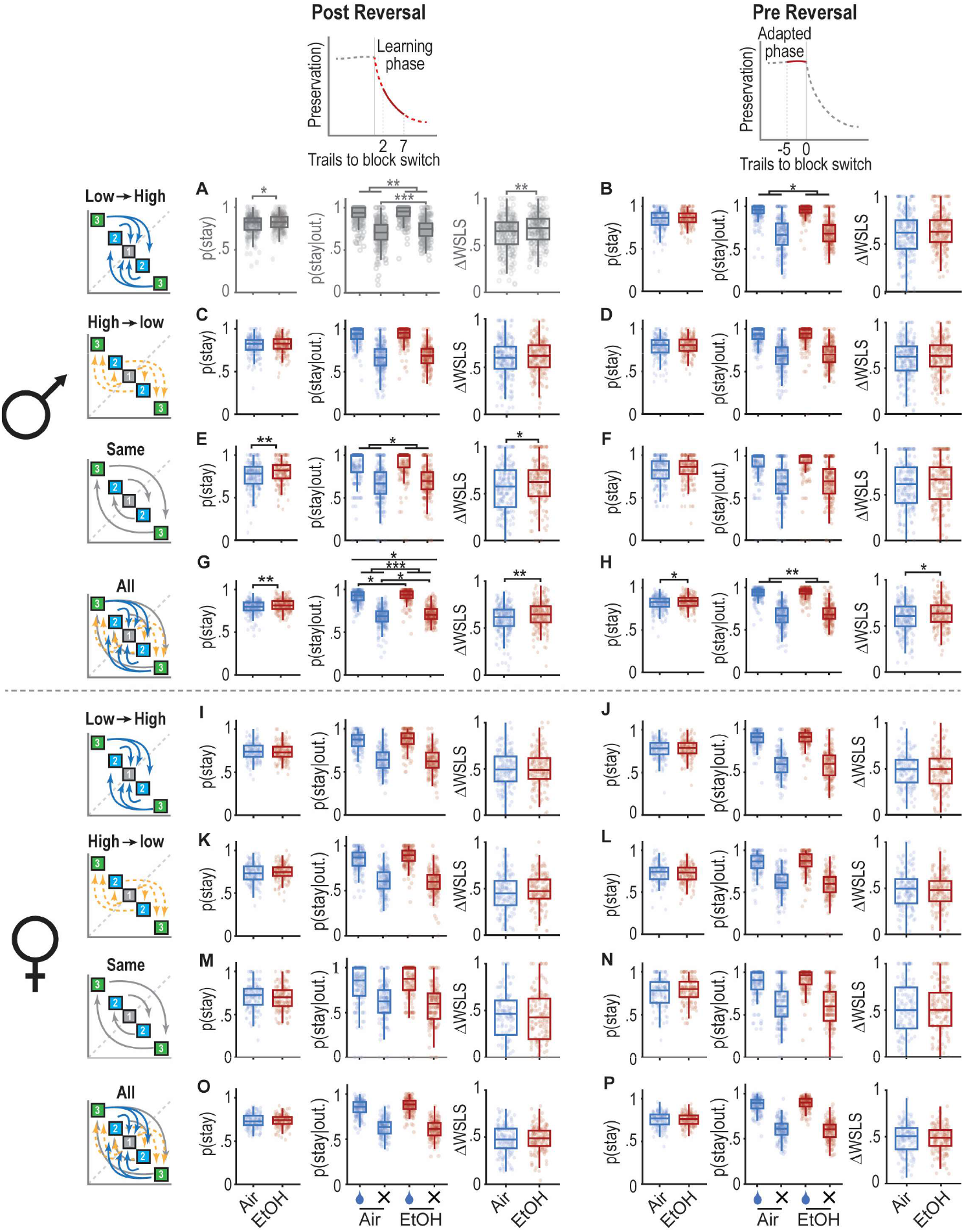
Stay probabilities in pre- and post-reversal under various cognitive challenge conditions. (**A**-**H**) Stay probability (left panels), Stay probability given reward (water drop icon) or no reward (cross, X, symbol) on the last trial (middle panels), and difference of win-stay lose-switch probability (ΔWSLS; right panels) in the post-reversal (A, C, E, and G) and pre-reversal (B, D, F, and H) phases during various uncertainty conditions in males. Data in A is duplicated from Figures 2H-2J. (**I**-**P**) Stay probability (left panels), conditional stay probability (middle panels), and difference of win-stay lose-switch probability (ΔWSLS; right panels) in the pre-reversal (I, K, M, and O) and post-reversal (J, L, N, and P) phases during various uncertainty conditions in females. Dots represent individual sessions in air and EtOH groups. N = 241 sessions from 9 male air rats and 229 sessions from 8 male EtOH rats.

**Figure S7.**
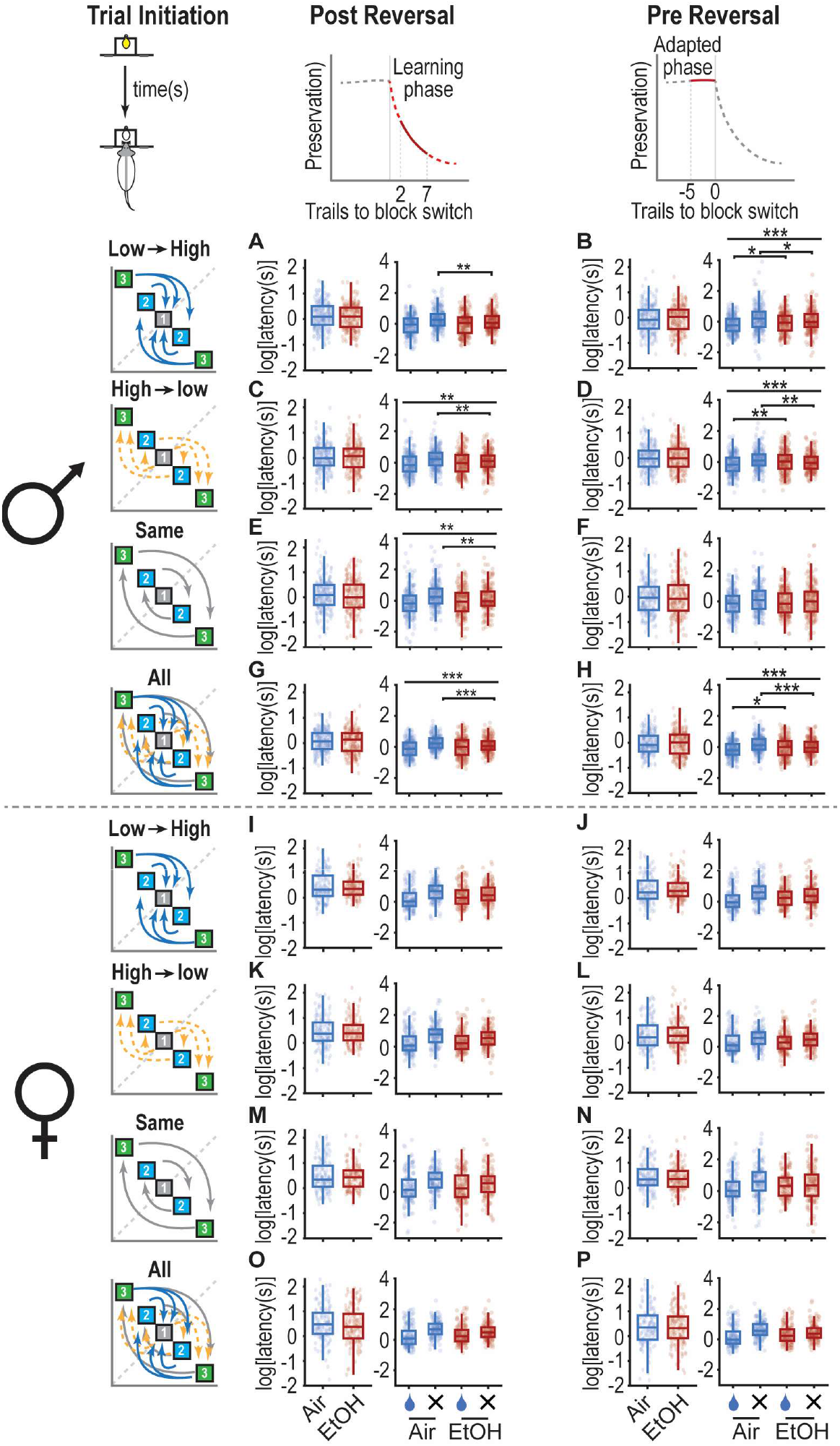
Latencies of trial initiation in pre- and post-reversal under various cognitive challenge conditions. (**A**-**H**) Latency of trial initiation (left panels) and latency of trial initiation conditioned on last trial outcomes (right panels) in the post-reversal (A, C, E, and G) and pre-reversal phases (B, D, F, and H) under various transitions in males. (**I**-**P**) Latency of trial initiation (left panels) and latency of trial initiation conditioned on last trial outcomes (right panels) in the post-reversal (A, C, E, and G) and pre-reversal phases (B, D, F, and H) under various transitions in females. Dots represent individual sessions in air and EtOH groups. N = 241 sessions from 9 male air rats and 229 sessions from 8 male EtOH rats.

**Figure S8.**
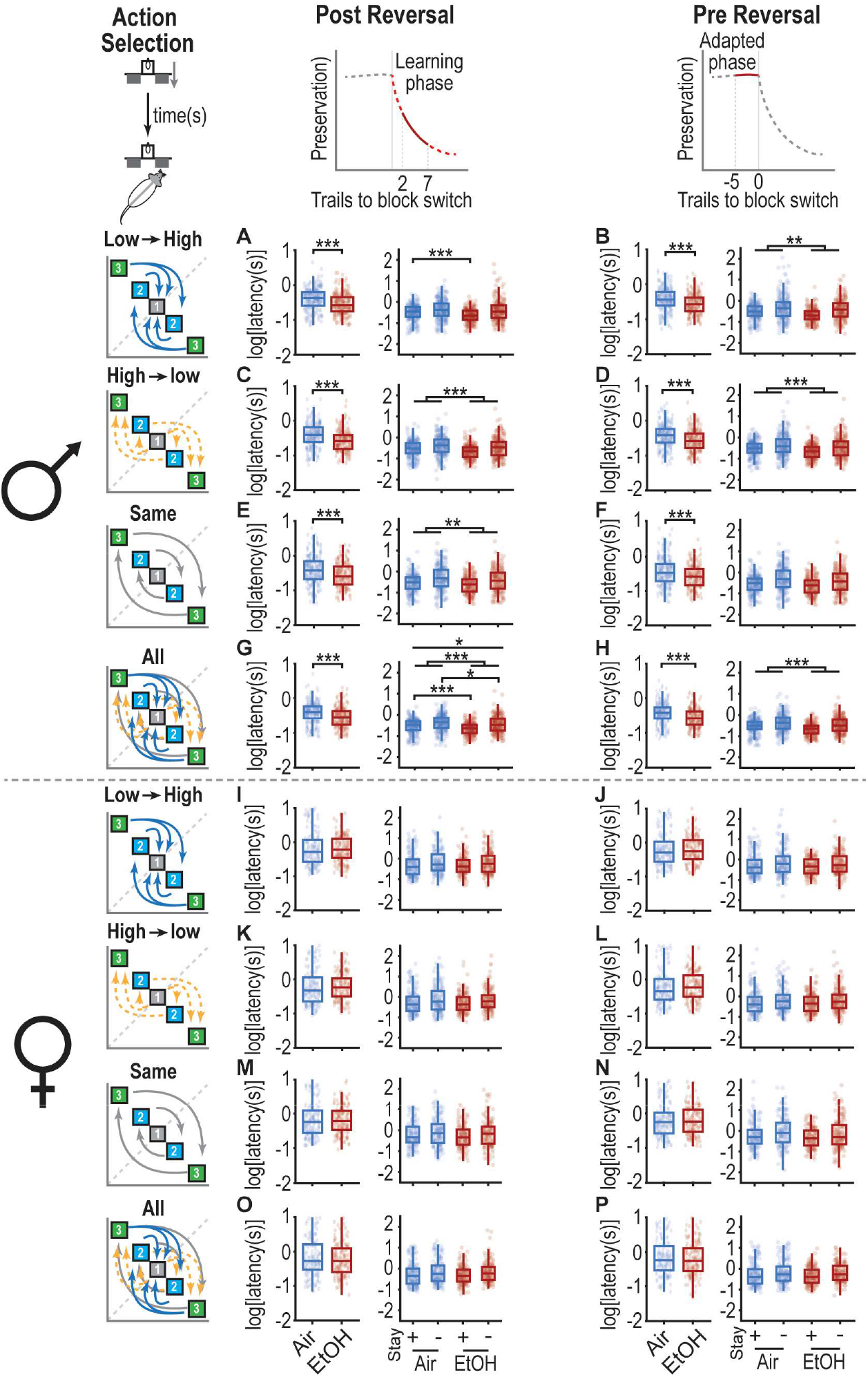
Latencies of action selection in pre- and post-reversal under various cognitive challenge conditions. (**A**-**H**) Latency of action selection (left panels) and latency of trial initiation conditioned on last trial outcomes (right panels) in the post-reversal (A, C, E, and G) and pre-reversal phases (B, D, F, and H) under various transitions in males. (**I**-**P**) Latency of trial initiation (left) and latency of trial initiation conditioned on last trial outcomes (right) in the post-reversal (A, C, E, and G) and pre-reversal phases (B, D, F, and H) under various transitions in females. Dots represent individual sessions in air and EtOH groups. N = 241 sessions from 9 male air rats and 229 sessions from 8 male EtOH rats.

**Figure S9.**
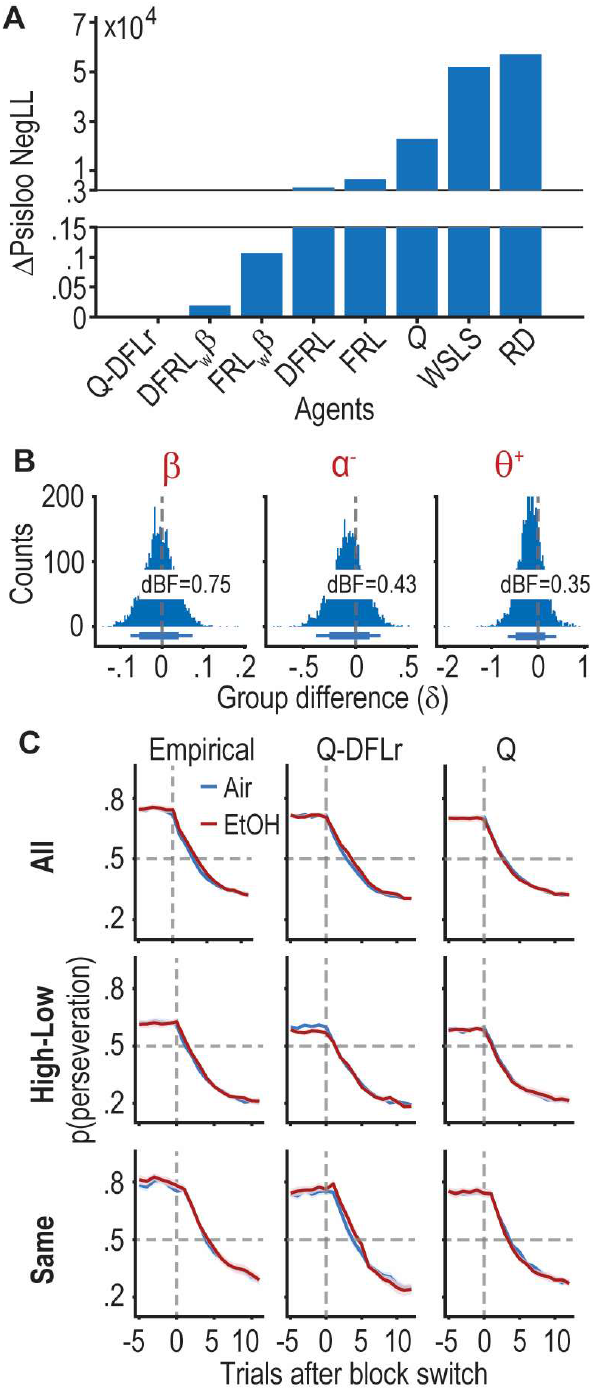
Reinforcement learning models fit trial-by-trial data from the dynaPRL task. (**A**) The bar plot shows the difference of Pareto-smoothed importance sampling (PSIS) with leave-one-out (LOO) negative likelihood (ΔPsisloo NegLL) between candidate models and the best fitting model, Q learning model with differential learning rate (Q-DFLr), on the most left on the x-axis. (**B**) HMC-sampled posterior densities of group difference, δ, representing differences between EtOH- and air-exposed rats in inverse temperature (β in the left panel), negative feedback-mediated learning rates (α^−^ in the middle panel), and positive feedback-mediated forgetting rates (θ^+^ in the right panel), across group-level hyperparameters. Bottom horizontal lines represent the 80% and 95% highest density interval (HDI). dBF: directed Bayes factor. (**C**) Choice probability of the initially preferred lever in all, High-low, and Same transitions. Data on the left panel are empirical (reproduced from Figure 1G), while the right panel depicts simulated data using the best-fitting parameters.

**Figure S10.**
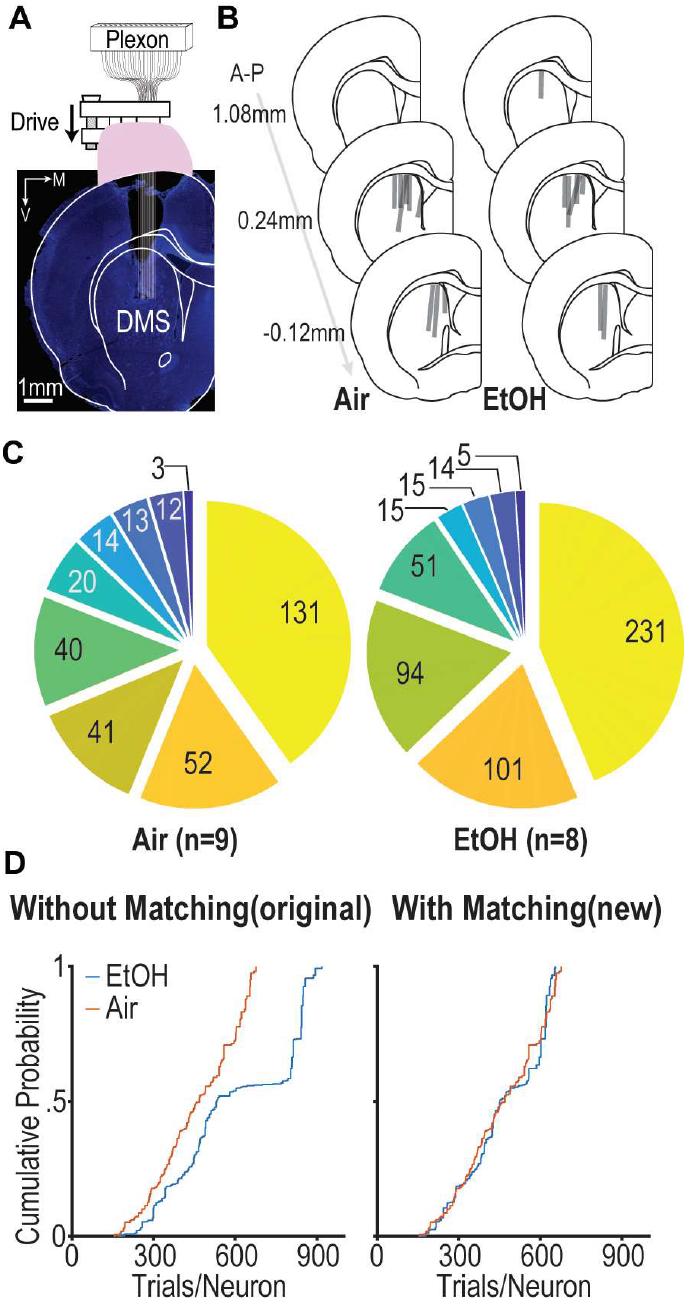
Single unit recording in the dorsomedial striatum (DMS). (**A**) The diagram illustrates a drivable electrode consisting of 16 tungsten wires connected to a Plexon headstage, which is implanted in the Dorsomedial Striatum (DMS). (**B**) A visualization of electrode placement and the range of recording across sessions for each animal, denoted by grey lines. (**C**) Pie charts show the aggregate count of neurons recorded from each rat. (**D**) The cumulative probability distribution shows the number of trials for each recorded neuron before (left) and after (right) trial-matching, which is conducted between EtOH and air groups.

**Figure S11.**
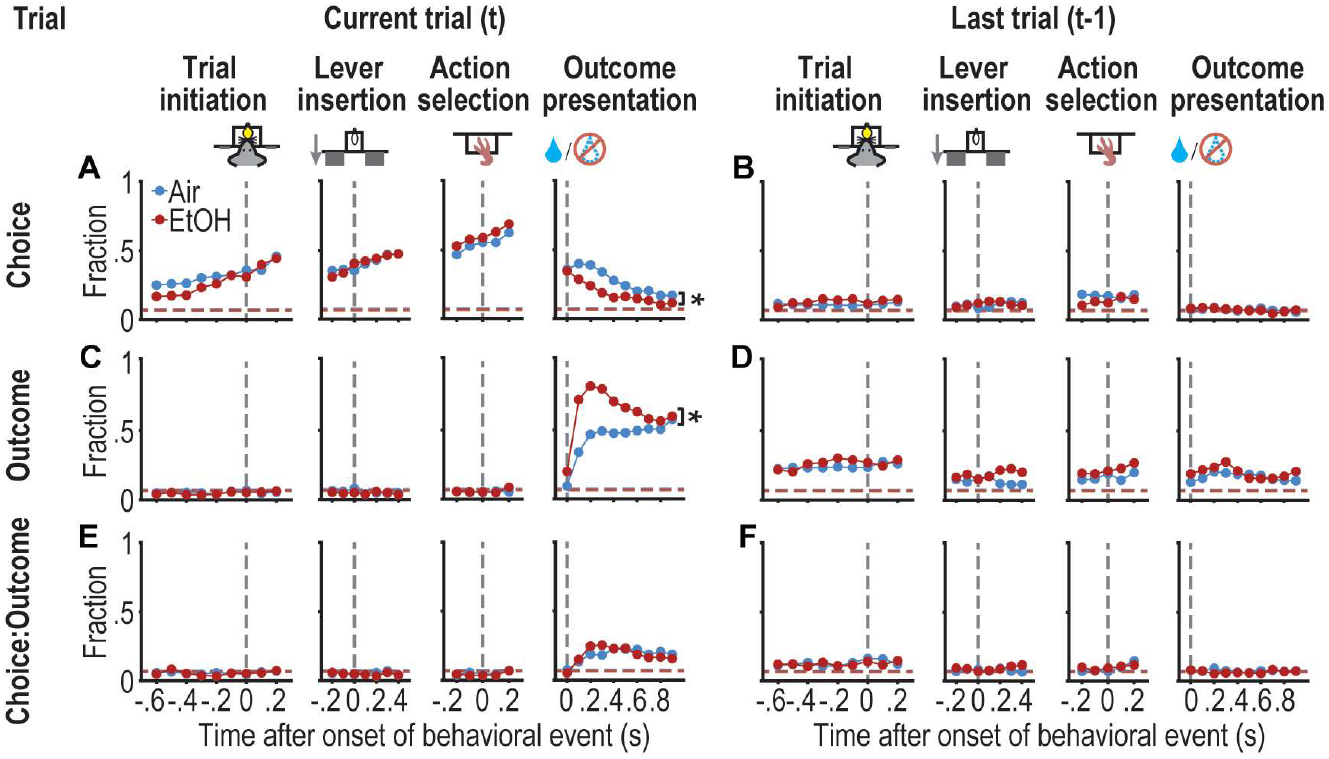
EtOH pre-exposure mainly alters the striatal choice and outcome signals in current trials. (**A**-**F**) Fraction of neurons exhibiting significant modulation by choice (A and B), outcome (C and D), and the interaction of choice and outcome (E and F) in current trial and last trial around behavioral events such as trials initiation, lever insertion, lever press, and outcome report. Vertical grey dashed lines indicate the onset of each behavior event. Horizontal blue and red dashed lines indicate the chance level for air and EtOH groups, respectively. N = 326 units from 9 male air rats, 523 units from 8 male EtOH rats.

**Figure S12.**
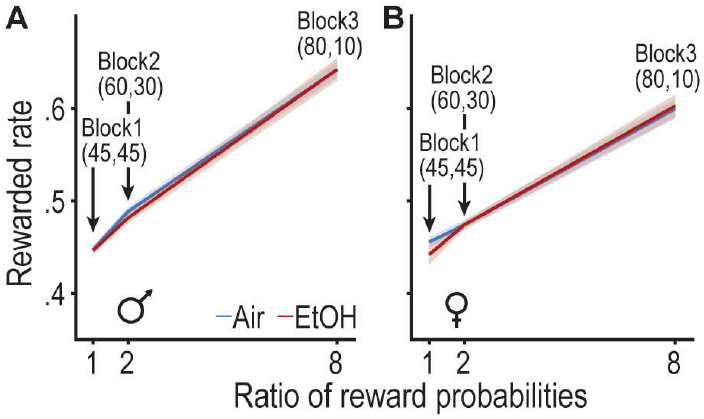
Reward rate as an indicator of block identity. (**A**-**B**) Reward rates (ratio of rewarded trials to total trials) linearly increase in correlation with the reward probability ratios, ranging from block 1 (45:45) to block 3 (80:10) for both males (A) and females (B). N = 13 EtOH rats (8 males, 5 females) and 14 air controls (9 males, 5 females).

**Figure S13.**
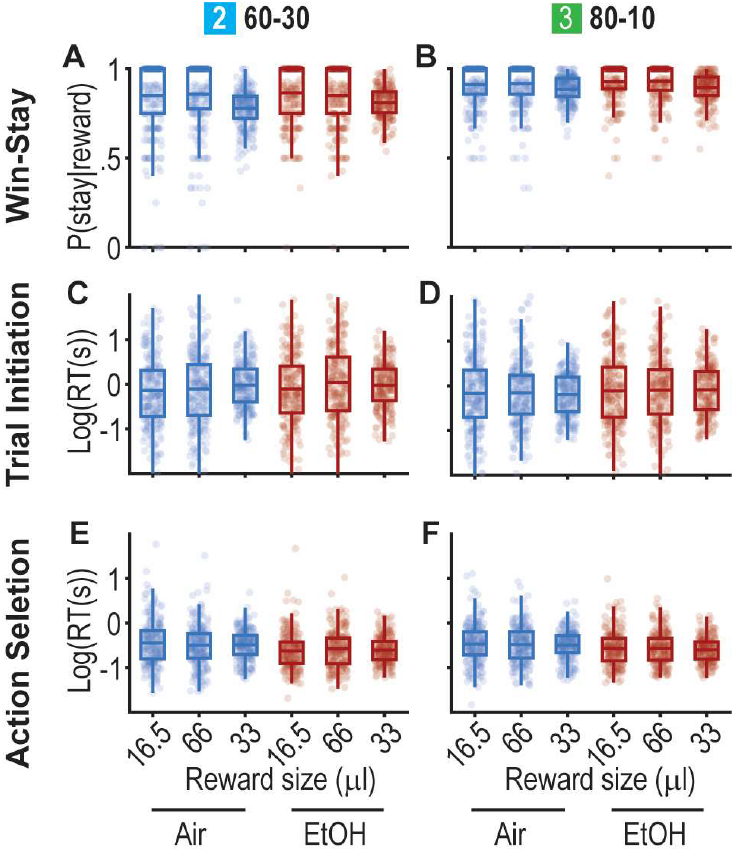
Rats do not respond to the size of the reward during the acquired phase in the dynaPRL task. (**A**-**F**) The size of the reward is altered with a 40% chance by doubling or halving the reward size (66 μl or 16.5 μl) compared to the 60% original size (33 μl) 12-trial after switching to block2 (45-45) and block3 (80-10). The probability of win-stay (A and B), latency of trial initiation (C and D), and action selection reaction time (E and F) are not significantly modulated by reward size. Dots represent the individual sessions.

## Notes

### Competing Interest Statement

The authors have declared no competing interest.

### Summary of Updates

This was the initial submission of my journal submission

## References

1. Barker, J.M., Corbit, L.H., et al. Corticostriatal circuitry and habitual ethanol seeking. Alcohol 49, 817–824 (2015).

2. Corbit, L.H. & Janak, P.H. Habitual Alcohol Seeking: Neural Bases and Possible Relations to Alcohol Use Disorders. Alcohol Clin Exp Res 40, 1380–1389 (2016).

3. Renteria, R., Baltz, E.T. & Gremel, C.M. Chronic alcohol exposure disrupts top-down control over basal ganglia action selection to produce habits. Nat Commun 9, 211 (2018).

4. Sjoerds, Z., de Wit, S., et al. Behavioral and neuroimaging evidence for overreliance on habit learning in alcohol-dependent patients. Transl Psychiatry 3, e337 (2013).

5. Sebold, M., Deserno, L., et al. Model-based and model-free decisions in alcohol dependence. Neuropsychobiology 70, 122–131 (2014).

6. Shields, C.N. & Gremel, C.M. Prior Chronic Alcohol Exposure Enhances Pavlovian-to-Instrumental Transfer. Alcohol (2021).

7. Corbit, L.H., Nie, H. & Janak, P.H. Habitual responding for alcohol depends upon both AMPA and D2 receptor signaling in the dorsolateral striatum. Front Behav Neurosci 8, 301 (2014).

8. Groman, S.M., Hillmer, A.T., et al. Midbrain D3 Receptor Availability Predicts Escalation in Cocaine Self-administration. Biol Psychiatry 88, 767–776 (2020).

9. Istin, M., Thiriet, N. & Solinas, M. Behavioral flexibility predicts increased ability to resist excessive methamphetamine self-administration. Addict Biol 22, 958–966 (2017).

10. Villiamma, P., Casby, J. & Groman, S.M. Adolescent reinforcement-learning trajectories predict cocaine-taking behaviors in adult male and female rats. Psychopharmacology 239, 2885–2901 (2022).

11. Turyabahika-Thyen, K. & Wolffgramm, J. Loss of flexibility in alcohol-taking rats: promoting factors. Eur Addict Res 12, 210–221 (2006).

12. Rodberg, E.M. & Vazey, E.M. Individual differences in behavioral flexibility predict future volitional ethanol consumption in mice. Alcohol 101, 37–43 (2022).

13. De Falco, E., White, S.M., et al. Impaired cognitive flexibility and heightened urgency are associated with increased alcohol consumption in rodent models of excessive drinking. Addict Biol 26, e13004 (2021).

14. Shnitko, T.A., Gonzales, S.W. & Grant, K.A. Low cognitive flexibility as a risk for heavy alcohol drinking in non-human primates. Alcohol 74, 95–104 (2019).

15. Czapla, M., Simon, J.J., et al. The impact of cognitive impairment and impulsivity on relapse of alcohol-dependent patients: implications for psychotherapeutic treatment. Addict Biol 21, 873–884 (2016).

16. Maillard, A., Laniepce, A., et al. Prognostic factors for low-risk drinking and relapse in alcohol use disorder: A multimodal analysis. Addict Biol 27 (2022).

17. Izquierdo, A. & Jentsch, J.D. Reversal learning as a measure of impulsive and compulsive behavior in addictions. Psychopharmacology 219, 607–620 (2012).

18. Ghods-Sharifi, S., Haluk, D.M. & Floresco, S.B. Differential effects of inactivation of the orbitofrontal cortex on strategy set-shifting and reversal learning. Neurobiol. Learn. Mem. 89, 567–573 (2008).

19. Reiter, A.M., Deserno, L., et al. Behavioral and Neural Signatures of Reduced Updating of Alternative Options in Alcohol-Dependent Patients during Flexible Decision-Making. J Neurosci 36, 10935–10948 (2016).

20. Chanraud, S., Reynaud, M., et al. Diffusion Tensor Tractography in Mesencephalic Bundles: Relation to Mental Flexibility in Detoxified Alcohol-Dependent Subjects. Neuropsychopharmacology : official publication of the American College of Neuropsychopharmacology 34, 1223–1232 (2009).

21. Jokisch, D., Roser, P., Juckel, G., Daum, I. & Bellebaum, C. Impairments in learning by monetary rewards and alcohol-associated rewards in detoxified alcoholic patients. Alcohol Clin Exp Res 38, 1947–1954 (2014).

22. Mason, B.J. Emerging pharmacotherapies for alcohol use disorder. Neuropharmacology 122, 244–253 (2017).

23. Macht, V., Elchert, N. & Crews, F. Adolescent Alcohol Exposure Produces Protracted Cognitive-Behavioral Impairments in Adult Male and Female Rats. Brain Sciences 10, 785 (2020).

24. Dannenhoffer, C.A., Robertson, M.M., et al. Chronic alcohol exposure during critical developmental periods differentially impacts persistence of deficits in cognitive flexibility and related circuitry. International Review of Neurobiology 160, 117–173 (2021).

25. Badanich, K.A., Becker, H.C. & Woodward, J.J. Effects of chronic intermittent ethanol exposure on orbitofrontal and medial prefrontal cortex-dependent behaviors in mice. Behav Neurosci 125, 879–891 (2011).

26. Kroener, S., Mulholland, P.J., et al. Chronic Alcohol Exposure Alters Behavioral and Synaptic Plasticity of the Rodent Prefrontal Cortex. PloS one 7, e37541 (2012).

27. Kuzmin, A., Liljequist, S., et al. Repeated moderate-dose ethanol bouts impair cognitive function in Wistar rats. Addict Biol 17, 132–140 (2012).

28. Ho, A.M.-C., Peyton, M.P., et al. Chronic Intermittent Ethanol Exposure Alters Behavioral Flexibility in Aged Rats Compared to Adult Rats and Modifies Protein and Protein Pathways Related to Alzheimer’s Disease. ACS Omega 7, 46260–46276 (2022).

29. Pickens, C.L., Gallo, M., Fisher, H., Pajser, A. & Ray, M.H. Alcohol Consumption during Adulthood Does Not Impair Later Go/No-Go Reversal Learning in Male Rats. NeuroSci 2, 166–176 (2021).

30. Aguirre, C.G., Stolyarova, A., et al. Sex-dependent effects of chronic intermittent voluntary alcohol consumption on attentional, not motivational, measures during probabilistic learning and reversal. PloS one 15, e0234729 (2020).

31. Ray, M.H., Hite, T., Gallo, M. & Pickens, C.L. Operant over-responding is more sensitive than reversal learning for revealing behavioral changes after withdrawal from alcohol consumption. Physiol Behav 196, 176–184 (2018).

32. Kim, H., Sul, J.H., Huh, N., Lee, D. & Jung, M.W. Role of striatum in updating values of chosen actions. J Neurosci 29, 14701–14712 (2009).

33. Izquierdo, A., Brigman, J.L., Radke, A.K., Rudebeck, P.H. & Holmes, A. The neural basis of reversal learning: An updated perspective. Neuroscience 345, 12–26 (2017).

34. Wang, J., Cheng, Y., et al. Alcohol elicits functional and structural plasticity selectively in dopamine D1 receptor-expressing neurons of the dorsomedial striatum. J Neurosci 35, 11634–11643 (2015).

35. Cheng, Y., Huang, C.C.Y., et al. Distinct synaptic strengthening of the striatal direct and indirect pathways drives alcohol consumption. Biol Psychiatry 81, 918–929 (2017).

36. Cheng, Y., Wang, X., et al. Prenatal Exposure to Alcohol Induces Functional and Structural Plasticity in Dopamine D1 Receptor-Expressing Neurons of the Dorsomedial Striatum. Alcohol Clin Exp Res 42, 1493–1502 (2018).

37. Cheng, Y., Xie, X., et al. Optogenetic induction of orbitostriatal long-term potentiation in the dorsomedial striatum elicits a persistent reduction of alcohol-seeking behavior in rats. Neuropharmacology 191, 108560 (2021).

38. Ma, T., Cheng, Y., et al. Bidirectional and long-lasting control of alcohol-seeking behavior by corticostriatal LTP and LTD. Nat Neurosci 21, 373–383 (2018).

39. Lu, J., Cheng, Y., et al. Alcohol intake enhances glutamatergic transmission from D2 receptor-expressing afferents onto D1 receptor-expressing medium spiny neurons in the dorsomedial striatum. Neuropsychopharmacology : official publication of the American College of Neuropsychopharmacology 44, 1123–1131 (2019).

40. Ma, T., Huang, Z., et al. Chronic alcohol drinking persistently suppresses thalamostriatal excitation of cholinergic neurons to impair cognitive flexibility. Journal of Clinical Investigation 132 (2021).

41. Tai, L.H., Lee, A.M., Benavidez, N., Bonci, A. & Wilbrecht, L. Transient stimulation of distinct subpopulations of striatal neurons mimics changes in action value. Nat Neurosci 15, 1281–1289 (2012).

42. Parker, N.F., Cameron, C.M., et al. Reward and choice encoding in terminals of midbrain dopamine neurons depends on striatal target. Nat Neurosci 19, 845–854 (2016).

43. Trantham-Davidson, H., Burnett, E.J., et al. Chronic alcohol disrupts dopamine receptor activity and the cognitive function of the medial prefrontal cortex. J Neurosci 34, 3706–3718 (2014).

44. Gilpin, N.W., Smith, A.D., et al. Operant behavior and alcohol levels in blood and brain of alcohol-dependent rats. Alcohol Clin Exp Res 33, 2113–2123 (2009).

45. Bi, J., Sun, J., Wu, Y., Tennen, H. & Armeli, S. A machine learning approach to college drinking prediction and risk factor identification. ACM Transactions on Intelligent Systems and Technology 4, 1–24 (2013).

46. Whelan, R., Watts, R., et al. Neuropsychosocial profiles of current and future adolescent alcohol misusers. Nature 512, 185–189 (2014).

47. Kinreich, S., Meyers, J.L., et al. Predicting risk for Alcohol Use Disorder using longitudinal data with multimodal biomarkers and family history: a machine learning study. Mol. Psychiatry 26, 1133–1141 (2021).

48. Grossman, C.D., Bari, B.A. & Cohen, J.Y. Serotonin neurons modulate learning rate through uncertainty. Curr Biol 32, 586–599 e587 (2021).

49. Funamizu, A., Ito, M., Doya, K., Kanzaki, R. & Takahashi, H. Uncertainty in action-value estimation affects both action choice and learning rate of the choice behaviors of rats. European Journal of Neuroscience 35, 1180–1189 (2012).

50. Trepka, E., Spitmaan, M., et al. Entropy-based metrics for predicting choice behavior based on local response to reward. Nat Commun 12, 6567 (2021).

51. Iigaya, K., Ahmadian, Y., et al. Deviation from the matching law reflects an optimal strategy involving learning over multiple timescales. Nat Commun 10 (2019).

52. Shin, E.J., Jang, Y., et al. Robust and distributed neural representation of action values. eLife 10 (2021).

53. Ridenour, T.A., Cottler, L.B., Compton, W.M., Spitznagel, E.L. & Cunningham-Williams, R.M. Is there a progression from abuse disorders to dependence disorders? Addiction 98, 635–644 (2003).

54. Manning, V., Wanigaratne, S., et al. Changes in Neuropsychological Functioning during Alcohol Detoxification. Eur Addict Res 14, 226–233 (2008).

55. Loeber, S., Duka, T., et al. Effects of Repeated Withdrawal from Alcohol on Recovery of Cognitive Impairment under Abstinence and Rate of Relapse. Alcohol and alcoholism 45, 541–547 (2010).

56. Moselhy, H.F. FRONTAL LOBE CHANGES IN ALCOHOLISM: A REVIEW OF THE LITERATURE. Alcohol and alcoholism 36, 357–368 (2001).

57. Chanraud, S., Martelli, C., et al. Brain Morphometry and Cognitive Performance in Detoxified Alcohol-Dependents with Preserved Psychosocial Functioning. Neuropsychopharmacology : official publication of the American College of Neuropsychopharmacology 32, 429–438 (2007).

58. Oscar-Berman, M., Shagrin, B., Evert, D.L. & Epstein, C. Impairments of brain and behavior: the neurological effects of alcohol. Alcohol Health Res World 21, 65–75 (1997).

59. Schulte, T., Müller-Oehring, E.M., Salo, R., Pfefferbaum, A. & Sullivan, E.V. Callosal involvement in a lateralized stroop task in alcoholic and healthy subjects. Neuropsychology 20, 727–736 (2006).

60. Pfefferbaum, A., Sullivan, E.V., et al. In vivo detection and functional correlates of white matter microstructural disruption in chronic alcoholism. Alcohol Clin Exp Res 24, 1214–1221 (2000).

61. Duka, T., Townshend, J.M., Collier, K. & Stephens, D.N. Impairment in cognitive functions after multiple detoxifications in alcoholic inpatients. Alcohol Clin Exp Res 27, 1563–1572 (2003).

62. Vanes, L.D., Holst, R.J., et al. Contingency Learning in Alcohol Dependence and Pathological Gambling: Learning and Unlearning Reward Contingencies. Alcohol Clin Exp Res 38, 1602–1610 (2014).

63. Corbit, L.H., Nie, H. & Janak, P.H. Habitual alcohol seeking: time course and the contribution of subregions of the dorsal striatum. Biol Psychiatry 72, 389–395 (2012).

64. Dominguez, G., Belzung, C., et al. Alcohol withdrawal induces long-lasting spatial working memory impairments: relationship with changes in corticosterone response in the prefrontal cortex. Addict Biol 22, 898–910 (2017).

65. Shnitko, T.A., Gonzales, S.W., Newman, N. & Grant, K.A. Behavioral Flexibility in Alcohol-Drinking Monkeys: The Morning After. Alcohol Clin Exp Res 44, 729–737 (2020).

66. Castillo-Carniglia, A., Keyes, K.M., Hasin, D.S. & Cerdá, M. Psychiatric comorbidities in alcohol use disorder. The Lancet Psychiatry 6, 1068–1080 (2019).

67. Birrell, J.M. & Brown, V.J. Medial Frontal Cortex Mediates Perceptual Attentional Set Shifting in the Rat. J Neurosci 20, 4320–4324 (2000).

68. Gangal, H., Xie, X., et al. Drug Reinforcement Impairs Cognitive Flexibility by Inhibiting Striatal Cholinergic Neurons. (Cold Spring Harbor Laboratory, 2022).

69. Bağci, B., Düsmez, S., et al. Computational analysis of probabilistic reversal learning deficits in male subjects with alcohol use disorder. Frontiers in psychiatry 13 (2022).

70. Morris, L.S., Baek, K., et al. Biases in the Explore–Exploit Tradeoff in Addictions: The Role of Avoidance of Uncertainty. Neuropsychopharmacology : official publication of the American College of Neuropsychopharmacology 41, 940–948 (2016).

71. Cockburn, J., Man, V., Cunningham, W.A. & O’Doherty, J.P. Novelty and uncertainty regulate the balance between exploration and exploitation through distinct mechanisms in the human brain. Neuron 110, 2691-2702.e2698 (2022).

72. Gershman, S.J. Uncertainty and Exploration. Decision (Wash D C) 6, 277–286 (2019).

73. Bechara, A., Dolan, S., et al. Decision-making deficits, linked to a dysfunctional ventromedial prefrontal cortex, revealed in alcohol and stimulant abusers. Neuropsychologia 39, 376–389 (2001).

74. Goudriaan, A.E., Oosterlaan, J., de Beurs, E. & van den Brink, W. Decision making in pathological gambling: A comparison between pathological gamblers, alcohol dependents, persons with Tourette syndrome, and normal controls. Cognitive Brain Research 23, 137–151 (2005).

75. Oglesby, M.E., Albanese, B.J., Chavarria, J. & Schmidt, N.B. Intolerance of Uncertainty in Relation to Motives for Alcohol Use. Cognitive Therapy and Research 39, 356–365 (2015).

76. Koob, G.F., Powell, P. & White, A. Addiction as a Coping Response: Hyperkatifeia, Deaths of Despair, and COVID-19. American Journal of Psychiatry 177, 1031–1037 (2020).

77. Oberlin, B.G. & Grahame, N.J. High-alcohol preferring mice are more impulsive than low-alcohol preferring mice as measured in the delay discounting task. Alcohol Clin Exp Res 33, 1294–1303 (2009).

78. Perkel, J.K., Bentzley, B.S., Andrzejewski, M.E. & Martinetti, M.P. Delay discounting for sucrose in alcohol-preferring and nonpreferring rats using a sipper tube within-sessions task. Alcohol Clin Exp Res 39, 232–238 (2015).

79. Smith, C.T., Steel, E.A., Parrish, M.H., Kelm, M.K. & Boettiger, C.A. Intertemporal Choice Behavior in Emerging Adults and Adults: Effects of Age Interact with Alcohol Use and Family History Status. Front Hum Neurosci 9, 627 (2015).

80. Barker, J.M., Bryant, K.G., Osborne, J.I. & Chandler, L.J. Age and Sex Interact to Mediate the Effects of Intermittent, High-Dose Ethanol Exposure on Behavioral Flexibility. Frontiers in pharmacology 8, 450 (2017).

81. Pratap, U.P., Patil, A., et al. Estrogen-induced neuroprotective and anti-inflammatory effects are dependent on the brain areas of middle-aged female rats. Brain Research Bulletin 124, 238–253 (2016).

82. Zhang, Q.-G., Wang, R., et al. Brain-derived estrogen exerts anti-inflammatory and neuroprotective actions in the rat hippocampus. Molecular and Cellular Endocrinology 389, 84–91 (2014).

83. Lei, H., Mochizuki, Y., et al. Sex difference in the weighting of expected uncertainty under chronic stress. Sci Rep 11, 8700 (2021).

84. Daw, N.D., Niv, Y. & Dayan, P. Actions, policies, values and the basal ganglia. in Recent Breakthroughs in Basal Ganglia Research (2005).

85. Groman, S.M., Thompson, S.L., Lee, D. & Taylor, J.R. Reinforcement learning detuned in addiction: integrative and translational approaches. Trends Neurosci 45, 96–105 (2022).

86. Beylergil, S.B., Beck, A., et al. Dorsolateral prefrontal cortex contributes to the impaired behavioral adaptation in alcohol dependence. Neuroimage Clin 15, 80–94 (2017).

87. Ito, M. & Doya, K. Validation of decision-making models and analysis of decision variables in the rat basal ganglia. J Neurosci 29, 9861–9874 (2009).

88. Samejima, K., Ueda, Y., Doya, K. & Kimura, M. Representation of action-specific reward values in the striatum. Science 310, 1337–1340 (2005).

89. Lau, B. & Glimcher, P.W. Value representations in the primate striatum during matching behavior. Neuron 58, 451–463 (2008).

90. Stalnaker, T.A., Calhoon, G.G., Ogawa, M., Roesch, M.R. & Schoenbaum, G. Neural correlates of stimulus-response and response-outcome associations in dorsolateral versus dorsomedial striatum. Front Integr Neurosci 4, 12 (2010).

91. Kim, H., Lee, D. & Jung, M.W. Signals for previous goal choice persist in the dorsomedial, but not dorsolateral striatum of rats. J Neurosci 33, 52–63 (2013).

92. Vandaele, Y., Mahajan, N.R., et al. Distinct recruitment of dorsomedial and dorsolateral striatum erodes with extended training. Elife 8 (2019).

93. Her, E.S., Huh, N., Kim, J. & Jung, M.W. Neuronal activity in dorsomedial and dorsolateral striatum under the requirement for temporal credit assignment. Sci Rep 6, 27056 (2016).

94. Bloem, B., Huda, R., et al. Multiplexed action-outcome representation by striatal striosome-matrix compartments detected with a mouse cost-benefit foraging task. Nat Commun 13 (2022).

95. Nishioka, T., Macpherson, T., Hamaguchi, K. & Hikida, T. Distinct Roles of Dopamine D1 and D2 Receptor-expressing Neurons in the Nucleus Accumbens for a Strategy Dependent Decision Making. (bioRxiv, 2021).

96. Bolkan, S.S., Stone, I.R., et al. Opponent control of behavior by dorsomedial striatal pathways depends on task demands and internal state. Nat Neurosci 25, 345–357 (2022).

97. Lee, J. & Sabatini, B.L. Striatal indirect pathway mediates exploration via collicular competition. Nature 599, 645–649 (2021).

98. Matamales, M., McGovern, A.E., et al. Local D2-to D1-neuron transmodulation updates goal-directed learning in the striatum. Science 367, 549–555 (2020).

99. Kwak, S. & Jung, M.W. Distinct roles of striatal direct and indirect pathways in value-based decision making. eLife 8, e46050 (2019).

100. O’Hare, J.K., Ade, K.K., et al. Pathway-Specific Striatal Substrates for Habitual Behavior. Neuron 89, 472–479 (2016).

101. Renteria, R., Cazares, C., et al. Mechanism for differential recruitment of orbitostriatal transmission during actions and outcomes following chronic alcohol exposure. eLife 10 (2021).

102. Stalnaker, T.A., Berg, B., Aujla, N. & Schoenbaum, G. Cholinergic Interneurons Use Orbitofrontal Input to Track Beliefs about Current State. J Neurosci 36, 6242–6257 (2016).

103. Hirokawa, J., Vaughan, A., Masset, P., Ott, T. & Kepecs, A. Frontal cortex neuron types categorically encode single decision variables. Nature 576, 446–451 (2019).

104. Schoenbaum, G. & Shaham, Y. The role of orbitofrontal cortex in drug addiction: a review of preclinical studies. Biol Psychiatry 63, 256–262 (2008).

105. Groman, S.M., Keistler, C., et al. Orbitofrontal Circuits Control Multiple Reinforcement-Learning Processes. Neuron 103, 734–746 e733 (2019).

106. Stalnaker, T.A., Liu, T.L., Takahashi, Y.K. & Schoenbaum, G. Orbitofrontal neurons signal reward predictions, not reward prediction errors. Neurobiol Learn Mem 153, 137–143 (2018).

107. Morris, R.W., Dezfouli, A., Griffiths, K.R., Le Pelley, M.E. & Balleine, B.W. The neural bases of action-outcome learning in humans. J Neurosci 42, JN-RM-1079–1021 (2022).

108. Alexander, W.H. & Brown, J.W. Medial prefrontal cortex as an action-outcome predictor. Nat Neurosci 14, 1338–1344 (2011).

109. Rushworth, M.F. & Behrens, T.E. Choice, uncertainty and value in prefrontal and cingulate cortex. Nat Neurosci 11, 389–397 (2008).

110. Hart, G., Bradfield, L.A. & Balleine, B.W. Prefrontal Corticostriatal Disconnection Blocks the Acquisition of Goal-Directed Action. J Neurosci 38, 1311–1322 (2018).

111. Koob, G.F. Alcoholism: Allostasis and Beyond. Alcohol Clin Exp Res 27, 232–243 (2003).

112. Jayaram-Lindström, N., Ericson, M., Steensland, P. & Jerlhag, E. Dopamine and Alcohol Dependence: From Bench to Clinic. in Recent Advances in Drug Addiction Research and Clinical Applications (2016).

113. Groman, S.M., Rich, K.M., Smith, N.J., Lee, D. & Taylor, J.R. Chronic exposure to methamphetamine disrupts reinforcement-based decision making in rats. Neuropsychopharmacology : official publication of the American College of Neuropsychopharmacology 43, 770–780 (2018).

114. Brown, V.M., Zhu, L., et al. Reinforcement Learning Disruptions in Individuals With Depression and Sensitivity to Symptom Change Following Cognitive Behavioral Therapy. JAMA Psychiatry 78, 1113–1122 (2021).

115. Wiehler, A., Chakroun, K. & Peters, J. Attenuated Directed Exploration during Reinforcement Learning in Gambling Disorder. J Neurosci 41, 2512–2522 (2021).

116. Kalhan, S., Schwartenbeck, P., Hester, R. & Garrido, M.I. People with a tobacco use disorder exhibit misaligned Bayesian belief updating by falsely attributing non-drug cues as worse predictors of positive outcomes compared to drug cues. Drug and Alcohol Dependence 256, 111109 (2024).

117. Vendruscolo, L.F. & Roberts, A.J. Operant alcohol self-administration in dependent rats: focus on the vapor model. Alcohol 48, 277–286 (2014).

118. Gilpin, N.W., Richardson, H.N., Cole, M. & Koob, G.F. Vapor inhalation of alcohol in rats. Curr Protoc Neurosci 9, 29 (2008).

119. Ottenheimer, D.J., Wang, K., et al. Reward activity in ventral pallidum tracks satiety-sensitive preference and drives choice behavior. Science Advances 6, eabc9321 (2020).

120. Ottenheimer, D.J., Bari, B.A., et al. A quantitative reward prediction error signal in the ventral pallidum. Nat Neurosci (2020).

121. Ottenheimer, D., Richard, J.M. & Janak, P.H. Ventral pallidum encodes relative reward value earlier and more robustly than nucleus accumbens. Nat Commun 9, 4350 (2018).

122. Meehan, C., Ebrahimian, J., Moore, W. & Meehan, S. Uniform Manifold Approximation and Projection (UMAP). (MATLAB Central File Exchange, https://www.mathworks.com/matlabcentral/fileexchange/71902, 2022).

123. Carpenter, B., Gelman, A., et al. Stan: A probabilistic programming language. Journal of statistical software 76 (2017).

124. Bowling, S.R., Khasawneh, M.T., Kaewkuekool, S. & Cho, B.R. A logistic approximation to the cumulative normal distribution. Journal of industrial engineering and management 2, 114–127 (2009).

125. Vehtari, A., Gelman, A. & Gabry, J. Practical Bayesian model evaluation using leave-one-out cross-validation and WAIC. Statistics and Computing 27, 1413–1432 (2017).

126. Elber-Dorozko, L. & Loewenstein, Y. Striatal action-value neurons reconsidered. Elife 7 (2018).

